# Weighing the DNA content of Adeno-Associated Virus vectors with zeptogram precision using nanomechanical resonators

**DOI:** 10.1101/2021.11.15.468734

**Authors:** Georgios Katsikis, Iris E. Hwang, Wade Wang, Vikas S. Bhat, Nicole L. McIntosh, Omair A. Karim, Bartlomiej J. Blus, Sha Sha, Vincent Agache, Jacqueline M. Wolfrum, Stacy L. Springs, Anthony J. Sinskey, Paul W. Barone, Richard D. Braatz, Scott R. Manalis

## Abstract

Quantifying the composition of viral vectors used in vaccine development and gene therapy is critical for assessing their functionality. Adeno-Associated Virus (AAV) vectors, which are the most widely used viral vectors for in-vivo gene therapy, are typically characterized using PCR, ELISA, and Analytical Ultracentrifugation which require laborious protocols or hours of turnaround time. Emerging methods such as Charge-Detection Mass Spectroscopy, Static Light Scattering, and Mass Photometry offer turnaround times of minutes for measuring AAV mass, but mostly require purified AAV-based reference materials for calibration. Here, we demonstrate a method for using Suspended Nanomechanical Resonators (SNR) to directly measure both AAV mass and aggregation from a few microliters of sample within minutes. We achieve a resolution near 10 zeptograms which corresponds to 1% of the genome holding capacity of the AAV capsid. Our results show the potential of our method for providing real-time quality control of viral vectors during biomanufacturing.

---

Adeno-Associated Viruses (AAV) are the most widely used viral vectors for in-vivo gene therapy due to their non-pathogenicity, low immunogenicity, and long-term gene expression^1^. However, AAV biomanufacturing is inefficient, producing only a small percentage (5–30%) of capsids containing the therapeutic gene; the majority of the produced capsids are empty which decreases the efficacy of gene therapy, increases its cost^2,3^, and has safety considerations^4^.

To increase the efficiency of AAV biomanufacturing, it is critical to provide real-time quality control to the process. As quality control is limited when using traditional molecular biology methods PCR and ELISA, in part due to their long turnaround times^5^, emerging methods based on mass measurement provide faster readouts for assessing the ratio of full (or ‘heavy’) to empty (or ‘light’) AAV capsids. In particular, mass spectrometers have been enhanced to independently measure mass through charge detection^6,7^, achieving attogram (or MDa) resolution. Multi-angle static light scattering detectors optically measure mass with sub-attogram (or kDa) resolution^8,9^. More recently, microscopy-based techniques correlating interferometric contrast and mass have also achieved kDa mass resolution^10–12^. Although these methods offer potential for real-time quality control, they depend on the optical or charge properties of a given AAV sample, and therefore mostly require purified AAV-based reference materials for calibration^5^.

Here, we directly measured AAV mass using mechanical resonators^13^. In particular, we used Suspended Nanochannel Resonators^14^ (SNR) to weigh AAVs in solution by measuring their buoyant mass, referred to from this point on as ‘mass’ (Figure 1a). The SNR is a hollow cantilever driven to vibrate at its resonant frequency *f*; when a single particle, such as a virus, flows through the cantilever, the resonant frequency of the cantilever transiently changes by *Δf*_*s*_ in proportion to the particle’s mass^15^.

**Figure 1.**
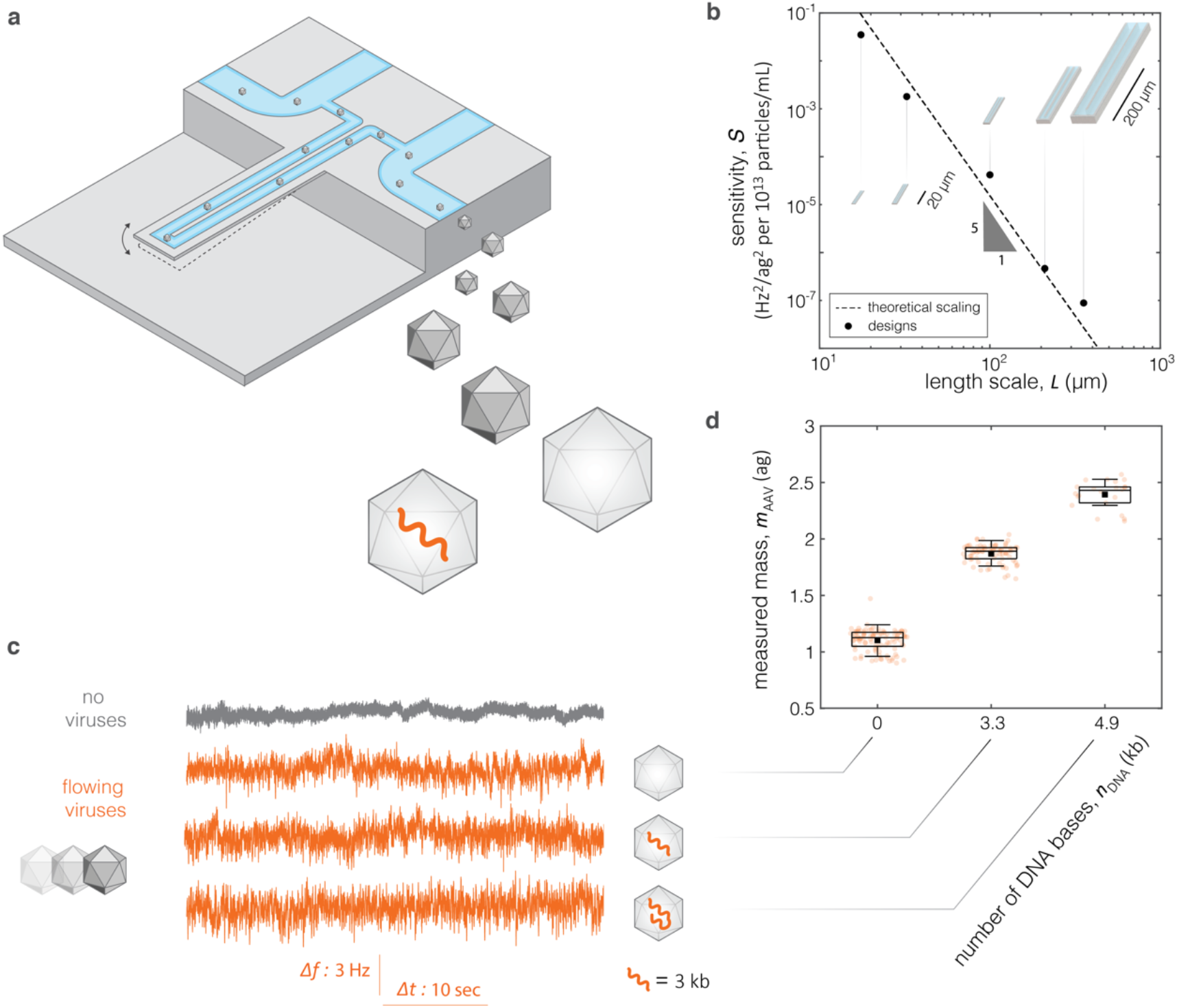
Concept of measuring mass of Adeno-Associated Viruses (AAV) in solution. **a**, Schematic of Suspended Nanochannel Resonator (SNR) featuring a hollow cantilever of length *L* = 17.5μ*m*, that vibrates at first resonant mode with frequency *f*. Inside the cantilever, we flowed solutions of AAVs with different DNA content or genetic constructs (denoted by orange color). **b**, Sensitivity *S* of root-mean-square *Δf*_*rms*_ of signal *Δf*(t) vs. length scale *L* of cantilever. Sensitivity is in units of Hz^2^ per nanoparticles of 1ag buoyant mass at a concentration of 10^13^ particle*s*/ml. Dashed line and triangle denote theoretical scaling with length *S*∼*L*^−5^, and black points represent experimental designs (Supplementary Note 4) **c**, Experimental time-series signals of change *Δf*(t) of resonant frequency exclusively due to noise in the absence of AAVs (gray), and due to flowing AAVs with distinct genetic constructs, having nominal number *n*_DNA_ of DNA kilobases (kb), *n*_DNA_ = 0, 3.3, 4.9 kb (from top to bottom in orange) **d**, Measured mass *m*_*AAV*_ of AAVs calculated from time-series data of panel **c** vs. *n*_DNA_. The central marks and black squares respectively indicate the median and mean. The bottom and top edges of the boxes respectively indicate the 25^th^ and 75^th^ percentiles. The bottom and top whiskers indicate the 5^th^ and 95^th^ percentiles. Percentiles are defined by assuming points follow normal distributions.

We previously used this approach to measure the mass of gold nanoparticles down to 10 nm in diameter^16^ corresponding to mass of 10 ag. However, a single AAV has a mass of 1 − 2 ag (molecular weight of *MW*_AAV_ = 3.8 − 5.4 MDa,^17,18^, Supporting Info S1), thus the frequency change *Δf*_*s*_ from a single AAV is occluded by noise. Although nanomechanical resonators have the potential to weigh single AAVs in a vacuum environment^19^, weighing AAVs in aqueous solution would enable rapid turnaround times which are ultimately required for real-time quality control.

To circumvent the noise limit for weighing single AAVs in solution, we flowed AAV samples at concentrations of 10^12^ − 10^13^ particles/ml such that tens to hundreds of AAVs simultaneously flow through the resonator (Figure 1a). This results in a complex time-series signal of frequency change *Δf*(t) which contains information about the mass of the particles as well as the characteristics of the flow. This concept has previously been used in microchannel resonators^20,21^ to measure the mass of polystyrene and gold nanoparticles weighing 20 ag, which is an order of magnitude heavier than AAVs. In addition, the signal *Δf*(*t*) also measures the volume of the particles^21^ in an approach similar to Dynamic Light Scattering^22^. Here, we scaled down the resonators from the micro-to nanoscale, and used spectral denoising to enable mass measurement of AAVs in the concentration range of 10^12^ − 10^13^ particle*s*/ml achieving a resolution near 10 zeptograms (zg) in a 10-minute sampling window.

To enable measurement of AAV mass, we first theoretically characterized the root-mean-square *Δf*_*rms*_ of signal *Δf*(t) as a function of the properties of the resonator in the form of a vibrating cantilever. It has been shown that *Δf*_*rms*_ (equivalent to the variance *σ*^2^ when the mean is *μ* = 0) is proportional to the product 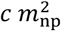 of average concentration *c* and average mass *m*_np_ of nanoparticles^20^. Here, we derived 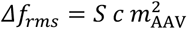 where *m*_*AAV*_ is the average mass of AAV nanoparticles, and the sensitivity 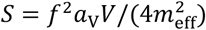 depends on the resonance frequency *f*, the volume *V* of the fluid channel, the effective mass *m*_eff_ of the cantilever^15^, and the volume utilization factor *a*_v_ related to the resonant mode *n*_m_ and the dimensions of the cantilever (Figure 1b, Supporting Info S2, S3, S4). The sensitivity *S*, for a length scale *L* of the cantilever, scales as *S*∼*L*^−5^ (Figure 1b, Supporting Info S2) indicating that the smaller the cantilever, the greater the *Δf*_*rms*_. Based on the scaling of *S* and the availability of previously characterized cantilevers^23^, we used the one with the smallest length *L* = 17.5 μ*m*, a fluid channel with a cross-sectional area 700 × 700 nm^2^, and a baseline resonant frequency *f* ≈ 4.5 MHz (Supporting Info S4).

Next, we measured AAV solutions with distinct genetic constructs, having nominal number *n*_DNA_ of DNA kilobases (kb), *n*_DNA_ = 0, 3.3 and 4.9 kb (Supporting Info S5). In the context of this study, we characterized the AAV5 serotype which is less prone to aggregation than AAV2 which is most commonly used in gene therapy^24^. Prior to testing, our AAV solutions were filtered and their purity was verified with SDS-PAGE and InstantBlue Staining (Supporting Information S5). When flowing these AAV samples in the SNR, we measured signals of frequency change *Δf*(t) which were visually distinct from the baseline noise (Figure 1c). Furthermore, we calculated the mass *m*_AAV_ from the signals *Δf*(t), and found that the measured mass for each AAV sample is distinguishable from one another (Figure 1d).

To validate our measurements, we performed experiments with nanoparticles of known mass as a reference standard for calibration. In particular, we used gold nanoparticles of nominal diameter *d*_Au,nom_ = 5 nm (Supporting Info S5). Using Dynamic Light Scattering we measured their mean hydrodynamic diameter and converted it to reference mass of *m*_Au,ref_ = 1.51 ag which is similar to that of AAV (Supporting Info S6). Furthermore, to gain insights into the experimental measurements, we developed a computational model based on the advection and diffusion of nanoparticles as they transit through the cantilever via a laminar flow, determined by a low 0 Reynolds number^25^ (Supplementary Info S7, Video S1). However, when we flowed gold nanoparticles in the cantilever, we found that calculating *Δf*_*rms*_ in the presence of noise overpredicts the nanoparticle mass (*m*_Au_ > 2 ag) in both the experiments and simulations (Figure 2, gray).

**Figure 2.**
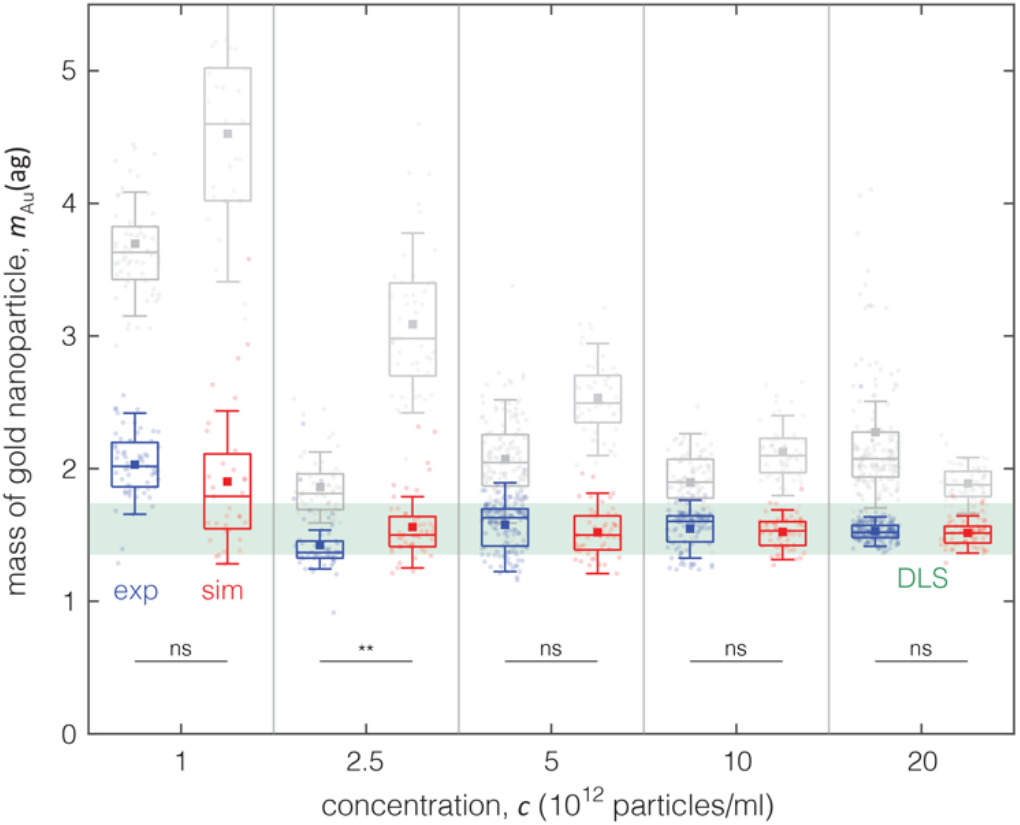
Validation of SNR measurements using nanoparticles of known mass. Mass of gold nanoparticles of nominal diameter *d*_*Au,nom*_ = 5 nm vs. concentration *c* (non-linear scale) specified with or without spectral denoising in experiments (exp: blue or gray) and simulations (sim: red or gray) (Supporting Info S9). When spectral denoising is used, the mass measurements from experiments and simulations are independent of concentration and consistent with the results from Dynamic Light Scattering (DLS, green band) revealing a mass of *m*_*Au,ref*_ = 1.51 ag corresponding to hydrodynamic diameter of *d* = 5.4 nm (Supporting Info S6). Box plots have similar notations as in Figure 1. The symbol * denotes *p* < 0.05, ** denotes *p* < 0.01 for the t-test, and *ns* denotes non-significant difference, between experiments and simulations.

To correct for the overprediction of nanoparticle mass, we first simulated the ‘pure’ signal *Δf*(t) caused by the flow of nanoparticles inside the cantilever in the absence of noise. We found that *Δf*(t) in the frequency domain is well-represented by a canonical Gaussian form 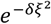, and when we calculated *Δf*_*rms*_ using a Gaussian fit in the frequency domain, we correctly predicted the nanoparticle mass (Supporting Info S8). We then experimentally characterized the noise in our system in the frequency domain and identified a canonical form 1/*ξ*^*a*^ of colored noise where *ξ* represents the spectral frequency, and the decay factor *a* lies in the range *a* = 1 − 2 (Supporting Info S9). Combining the canonical forms for ‘pure’ signal and noise, we developed a spectral denoising method that calculates *Δf*_*rms*_ in the frequency domain while neutralizing the effect of noise (Supporting Info S9). By applying spectral denoising to experiments and simulations of gold nanoparticles, we obtained results which are consistent with those obtained from Dynamic Light Scattering (Figure 2, blue and red). Remarkably, the spectral denoising method applied to data from both the experiments and simulations leads to a mass resolution of 10 − 100 zg (1 zg = 10^−3^ ag) in a 10-minute sampling window for concentrations of *c* = 5 − 20 × 10^12^ particles/ml (Supporting Info S10).

To validate our SNR measurements, we used established methods for characterizing AAVs^5,26^ (Supporting Information S5). Specifically, we characterized an AAV sample encompassing a genetic construct of Green Fluorescent Protein (GFP) with a nominal number *n*_DNA_ = 3.3 kb using Alkaline Agarose Gel Electrophoresis (AAGE) and Analytical Ultracentrifugation (AUC) (Figure 3a,b). AAGE revealed the presence of two distinct DNA bands; the ‘heavy’ (2.8 kb) is close to the nominal number of the construct, and the ‘heavy cs’ (4.7 kb) likely corresponds to a maximum-size construct that can be packaged inside the capsid as a result of an unintended formation of a self-complementary (cs) DNA sequence^27^ (Figure 3a). AUC and AAGE consistently revealed the presence of these two constructs. In addition, AUC identified residual DNA, ‘light’ (∼0 kb) and ‘intermediate’ (<2.8 kb) capsids as well as aggregates (Figure 3b, Supporting Info S11).

**Figure 3.**
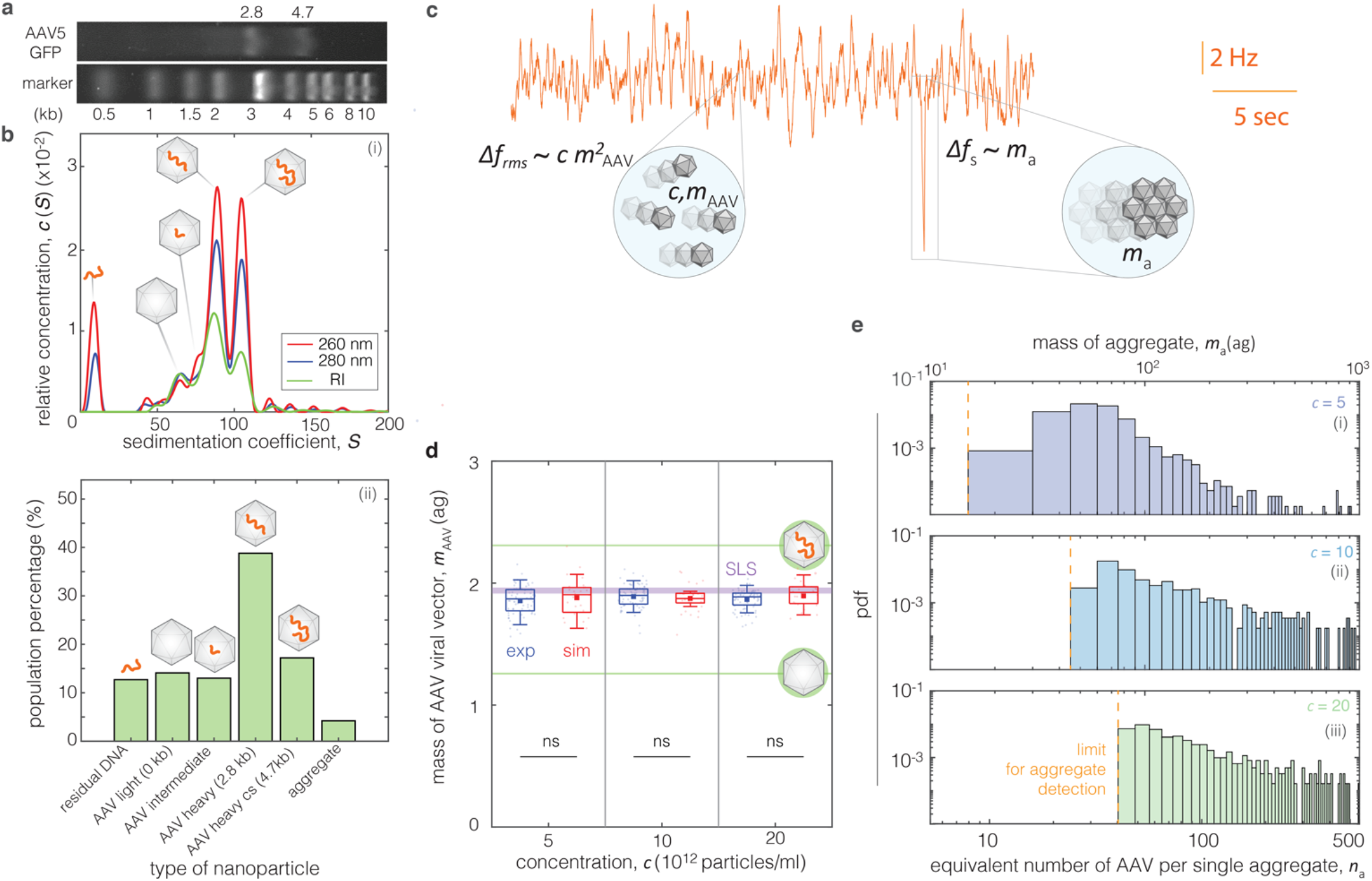
Measuring AAV mass and aggregation. **a**, Alkaline Gel Electrophoresis (AAEG) analysis of AAV-GFP (top) vs. marker (bottom) measuring number of DNA bases (kb) **b**, Analytical Ultracentrifugation (AUC) analysis of AAV-GFP using Refractive Index (RI) signal (green) and 260, 280 nm UV wavelengths (red, blue) (i). *c*_260_/*c*_280_ > 1 indicates AAV capsids filled with DNA^26^ (e.g., ‘heavy capsids’) Population percentage (ii) extracted from (i) with numbers corresponding to peaks indicating different types of AAV (insets). The letters ‘cs’ in ‘AAV heavy cs’ denote complementary strand, filling up the capsid to its maximum holding capacity (4.7 kb) **c**, Example of experimental signal *Δf*(t) showing the simultaneous measurements of the average AAV mass *m*_*AAV*_ of concentration *c* (*Δf*_*rms*_), and the mass *m*_*v*_ of individual aggregates (*Δf*_*s*_ < 0). Box plots have similar notation as in Figures 1,2. **d**, Mass *m*_*AAV*_ of AAV viral vector with genetic construct of GFP vs. concentration *c* (non-linear scale) measured using spectral denoising in both experiments (exp: blue) and simulations (sim: red). The purple band indicates the range of mass specified using Static Light Scattering (SLS) where its width is standard deviation from n = 3 experiments. The green bands correspond to the types of AAV from panel **b. e**, Probability density functions of mass *m*_*a*_ of aggregates (top horizontal axis), also expressed as equivalent number of AAVs per single aggregates (top horizontal axis) for three concentrations (i-iii). The limit for detection of aggregates (orange), is governed by *Δf*_*rms*_ being dependent on *c*.

We found that *Δf*_*rms*_ from SNR method (Figure 3d) 155 measures AAV mass that is consistent with the results from AAGE and AUC. By using spectral denoising (Supporting Info S9), simulations (Supporting Info S7) and conversions of measured AAV mass to DNA content (Supporting Info S1), our measurements are within the margin specified by AUC (Figure 3d). In addition, our results are consistent with Static Light Scattering (SLS) measurements, an orthogonal method recently used for characterizing AAVs^9^ (Figure 3d, Supporting Info S12). Overall, we found that our SNR measurements are reliable down to a concentration limit of *c* ≈ 5 × 10^12^ particle*s*/ml (Figure 3d). Below that limit, the contribution of noise to *Δf*_*rms*_ is higher than that of the AAV signal (Supporting Info S13). Notably, the resolution of AAV mass measurements, similar to that of gold nanoparticles (Figure 2), is maintained at a near 10 zg level at the upper concentration range *c* = 2 × 10^13^ particles/ml (Supporting Info S10).

Simultaneously with measuring AAV mass, we determined the mass *m*_a_ of single aggregates, each of which manifests as a transient decrease in resonant frequency (Figure 3c, *Δf*_*s*_ < 0) as each aggregate passes through the cantilever. As a basis for comparison, we confirmed the presence of soluble aggregates in our AAV sample using Dynamic Light Scattering (DLS) although DLS cannot in principle resolve the heterogeneity of aggregates (Supporting Information S6). The presence of aggregates was not only evident in Analytical Ultracentrifugation (Figure 3b) but also in chromatographic methods. In particular, Anion Exchange Chromatography analysis (AEX)^28^ exhibited long tailing at higher elution times (Supporting Information S14) while Size Exclusion Chromatography (SEC) showed long tailing at the lower elution times (Supporting Information S15). Using SNR, we directly measured the mass of these single aggregates and characterized their heterogeneity (Figure 3e). Importantly, the detection limit for weighing single aggregates by our method depends on the baseline frequency noise of *Δf*(t) which is essentially governed by *Δf*_*rms*_, being dependent on the AAV concentration (Supporting Info S8). However, within the limits of aggregate detection, we found that the mass heterogeneity of aggregates was similar for the three tested AAV concentrations in the range *c* = 5 − 20 × 10^12^ particle*s*/ml (Figure 3e).

We leveraged the measurement of AAV mass to convert our readout to a ratio of full to empty or ‘heavy to light’ capsids which, along with aggregation, is a critical quality attribute of a given AAV product.^5^ We thus measured the mass of AAV mixtures with different volume ratios of AAV ‘heavy’ as AAV-GFP (*n*_DNA_ = 3.3 kb) and AAV ‘light’ as AAV-empty without a nominal DNA construct (*n*_DNA_ = 0 kb). We observed that our experimental results are consistent with our simulations with a trend of increasing mass for mixtures of higher percentage of heavy-to-light capsids (Figure 4a). In addition, given the theoretical mass values for AAVs based on their DNA content, (Supporting Info S1), we converted the mass *m*_*AAV*_ to percentage of heavy AAV capsids, obtaining consistent percentages with those determined by Static Light Scattering for the same mixture ratios (Figure 4b).

**Figure 4.**
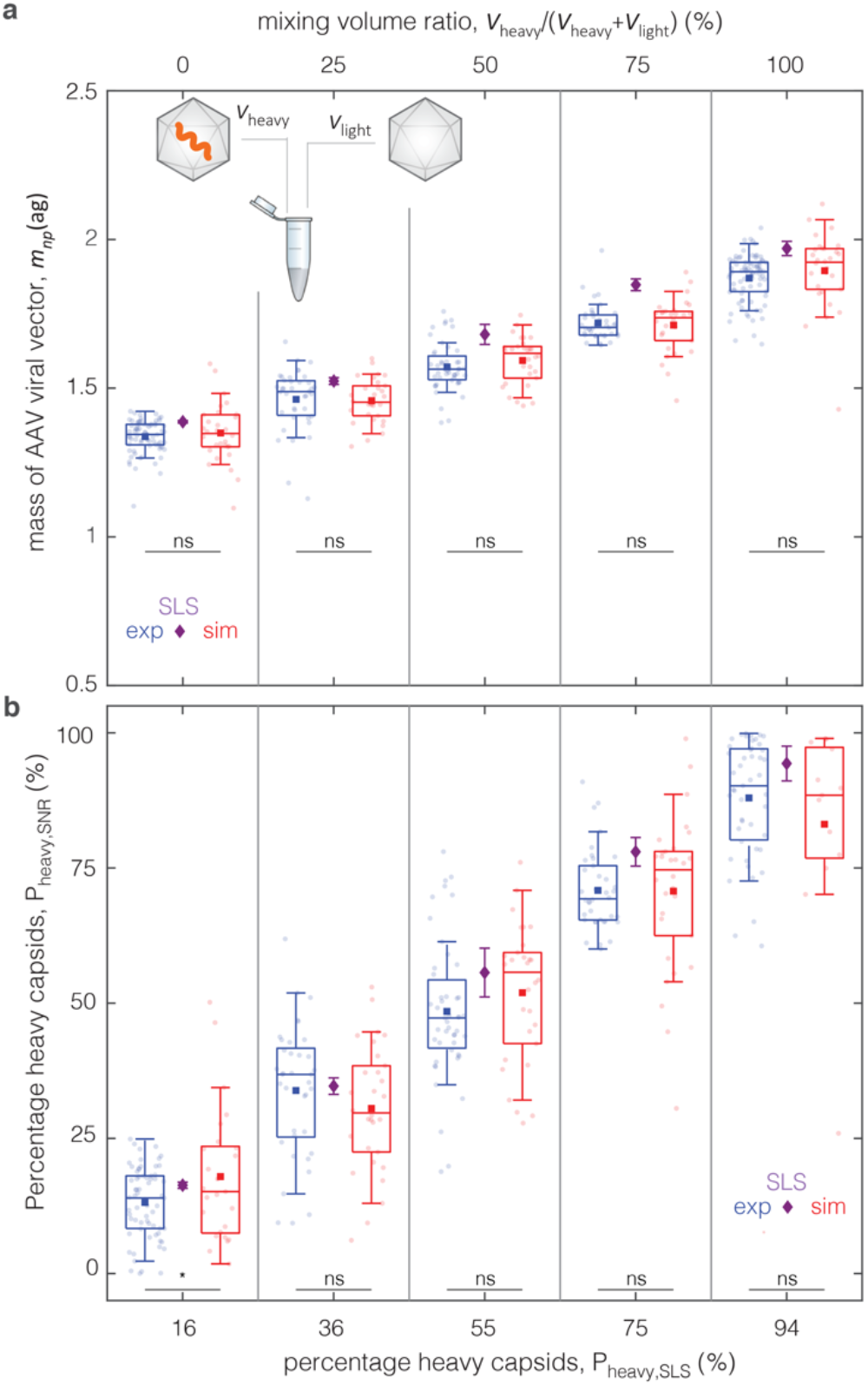
Calculation of percentage of heavy capsids using mass measurement of AAV. **a**, Mass *m*_*AAV*_ of mixtures of two types of AAV samples : i) GFP ‘heavy’ and ii) empty ‘light’ vs. mixing volume ratio for experiments (exp: blue) simulations (sim: red). **b**, Percentage ofheavy capsids, defined as *P*_*heavy*_ = (*m*_AAV_ − *m*_light_)/(*m*_*heavy*_ − *m*_light_) for SNR (vertical axis) vs SLS (horizontal axis) where *m*_light_, *m*_heavy_ are theoretical values for buoyant mass of AAV (Supporting Info S1) when they respectively contain no DNA (*n*_DNA_ = 0 kb) or DNA of the full genetic construct (*n*_DNA_ = 4.7 kb). The horizontal axis corresponds to the percentage of full capsids calculated using Static Light Scattering (SLS), also denoted by purple diamonds. Error bars denote standard deviation in SLS experiments (*n* = 3 per mixing volume ratio). Boxplots have similar notation as in Figures 1,2.

Our SNR results establish the foundations for a real-time method for characterizing AAV-based viral vectors in solution as an orthogonal method to Static Light Scattering. Although SNR cannot resolve heterogeneity of individual AAVs as opposed to charge-detection mass spectroscopy or mass photometry, it does not require purified, AAV-based reference materials for calibrating the mass measurement^7,12,29^, and has a turnaround time of 10 minutes. Importantly, our method enables simultaneous quantification of both the average DNA content and individual aggregates, which constitute two critical quality attributes of a given AAV product^5^. Furthermore, it is a flow-through method requiring minimal amount of sample (1 − 5 μL) when compared to analytical ultracentrifugation which is laborious, and typically requires more than 100-fold larger sample volumes (400 μL)^30^.

We envision that the success of AAV characterization will rely on a synergistic pipeline of multiple methods that provide different signal detection readouts. In addition to measurement of AAV mass and concentration (or ‘titer’), upstream purification is equally important as AAV measurements are prone to biases due to the presence of impurities^5^. In addition to SNR, chromatography-based methods, such as size exclusion or anion exchange chromatography remain indispensable elements of the overall pipeline of AAV characterization^5,9,31,32^ despite their limitations in resolving capsid heterogeneity. Such characterization coupled with molecular engineering, process development^33^, as well as mathematical modelling/simulation^34^ of the AAV biomanufacturing process may lead to new paradigms for scaling up production of viral vectors. Although our approach is showcased here for AAV5, we envision it is also applicable to a broader context of viral vectors, thereby paving the ground for increasing the efficacy of gene therapy treatments while maintaining their affordability and clinical safety.

## ASSOCIATED CONTENT

The supporting information is available free of charge at []

Detailed information of supporting figures, theory, and methods (PDF).

Video S1 showing simulation of Adeno-Associated viruses flowing through suspended nanochannel resonator (top), with resulting signal of change in resonant frequency *Δf*(t) as viruses are flowing through the resonator (bottom).

## Author Contributions

G.K. and S.R.M. conceived the study. G.K., W.W., V.S.B., and S.R.M. designed the research. G.K. carried out the theoretical analysis, and performed the experiments with the suspended nanochannel resonators. V.A. provided the suspended nanochannel resonators. I.E.H. developed the computational model. G.K. and I.E.H. performed the simulations. S.S. did initial PCR and ELISA measurements and supported the early development of the project. W.W., N.L.M., O.A.K., and B.J.B. performed the experiments using established methods for characterizing AAVs. G.K., I.E.H. N.L.M., O.A.K., B.J.B., and W.W. analyzed the results. J.M.W., S.L.S., A.J.S., P.W.B., V.A., R.D.B. and S.R.M. guided the work. G.K., I.E.H., S.R.M. wrote the paper with feedback from W.W, V.S.B., V.A., J.M.W., S.L.S., A.J.S., P.W.B, B.J.B. and R.D.B.

## Funding Sources

This study is supported by a grant from the US Food and Drug Administration (grant ID 1R01FD006584-02, Continuous Viral Vector Manufacturing based on Mechanistic Modeling and Novel Process Analytics). S.R.M. and G.K. acknowledge support from the Virginia and D.K. Ludwig Fund for Cancer Research.

## Notes

S.R.M. is a co-founder of Travera and Affinity Biosensors, which develops technologies relevant to the research presented in this work.

## ACKNOWLEDGMENT

We acknowledge Julie Sutton for discussion on noise characterization, and help with experimental configurations of the suspended nanochannel resonator. We acknowledge Elizabeth ‘Betsy’ Skrip from MIT Center for Biomedical Innovation for aesthetically supporting the design of figures.

