## Supporting Information for "Weighing the DNA content of Adeno-Associated Virus vectors with zeptogram precision using nanomechanical resonators"

[1] Koch Institute for Integrative Cancer Research, Massachusetts Institute of Technology, Cambridge, MA 02139 USA.
[2] BioMarin Pharmaceutical, Inc., Novato, CA, 94929, USA.
[3] BioMarin Pharmaceutical, Inc., San Rafael, CA, 94929, USA.
[4] Center for Biomedical Innovation, Massachusetts Institute of Technology, Cambridge, MA, 02139 USA.
[5] Université Grenoble Alpes, CEA, LETI, 38000, Grenoble France.
[6] Department of Biology, Massachusetts Institute of Technology, Cambridge, MA, 02139 USA.
[7] Department of Chemical Engineering, Massachusetts Institute of Technology, Cambridge, MA, 02139 USA.
[8] Department of Mechanical Engineering, Massachusetts Institute of Technology, Cambridge, MA, 02139 USA.
[9] Department of Biological Engineering, Massachusetts Institute of Technology, Cambridge, MA, 02139 USA.
[*].

### S1. Calculation of AAV buoyant mass and percentage of heavy capsids

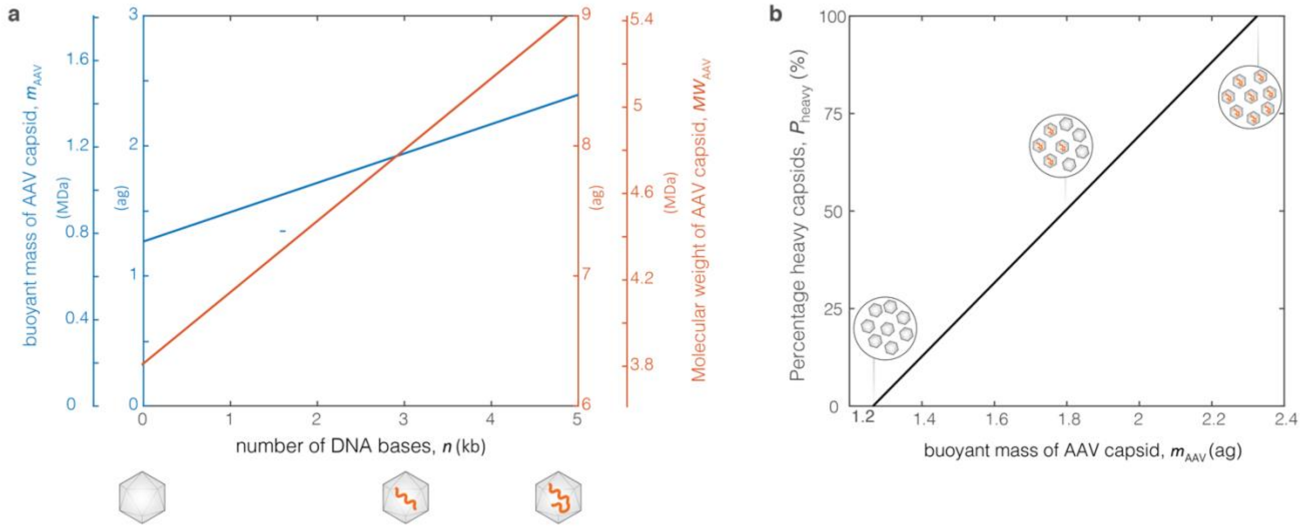


**Figure S1 | Calculation of buoyant mass and percentage of heavy capsids of Adeno-Associated Virus (AAV). a,** Calculation of buoyant mass $m_{\text{AAV}}$ of AAV capsid from molecular weight[13,14] $MW_{\text{AAV}}$ assuming fluid density $\rho = 1{,}000\ \text{kg/m}^3$, protein density $\rho_{\text{p}} = 1{,}250\ \text{kg/m}^3$, DNA density $\rho_{\text{DNA}} = 1{,}700\ \text{kg/m}^3$. The capsid of AAV serotype 5 used in this study consists of three types of viral proteins (VP) with $MW_{\text{VP1}} = 87$ kDa, $MW_{\text{VP2}} = 73$ kDa, $MW_{\text{VP3}} = 62$ kDa, and corresponding number ratios $1{:}\,1{:}\,10$ for a total number of 60 proteins. **b**, Percentage of heavy capsids defined as $P_{\text{heavy}} = (m_{\text{AAV}} - m_{\text{light}})/(m_{\text{heavy}} - m_{\text{light}})$, where $m_{\text{light}}, m_{\text{heavy}}$ are theoretical values for buoyant mass of AAV when they, respectively, are empty or 'light' containing no DNA ($n_{\text{DNA}} = 0$ kb), and are full or 'heavy' containing the DNA of the full genetic construct ($n_{\text{DNA}} = 4.7$ kb) from panel **a**.

### S2. Theory of Measurement

When a solution of AAV flows through a cantilever vibrating at resonant frequency $f$, it causes complex times-series signal of frequency change[18,19] $\Delta f(t)$ with a root mean square $\Delta f_{rms} = \overline{\Delta f^2(t)}$ value that is equal to:

$$\Delta f_{rms} = \frac{f^2 c\, m_{\mathrm{AAV}}^2}{4 m_{\mathrm{eff}}^2} \iiint_V W^4 dV \tag{1}$$

where $m_{\mathrm{AAV}}$ is the average buoyant mass of the AAV, $c$ is average concentration of AAV particles, $m_{\mathrm{eff}}$ is the effective mass of the cantilever, $W(x)$ is the shape of displacement of cantilever along its length $x$ for given resonant mode[12], and $V$ represents the volume of the fluid channel.

Since the cantilever is single-clamped on one side, the fluid channel has a U-shape for entry and exit of particles into the cantilever. The volume of the fluid channel is:

$$V = \underbrace{2\, H\, W L_{\mathrm{fluid}}}_{V_{\mathrm{main}}} + \underbrace{H\, W_{\mathrm{int}}\, (L_{\mathrm{fluid}} - L_{\mathrm{wall}})}_{V_{\mathrm{tip}}} \tag{2}$$

The variables $H, W, W_{\mathrm{int}}, L_{\mathrm{fluid}}, L_{\mathrm{wall}}$ in equation (2) refer to the dimensions of the cantilever (Supporting Info S4).

Since the shape of displacement $W(x)$ depends only on $x$, for each of the fluid volumes $V_{\mathrm{main}}$, $V_{\mathrm{tip}}$ we calculated the terms:

$$\iiint_{V_{\mathrm{main}}} W^4 dV = 2H\, W \int_{x=0}^{x=L_{\mathrm{fluid}}} W^4\, dx \tag{3}$$

$$\iiint_{V_{\mathrm{tip}}} W^4 dV = H\, W_{\mathrm{int}} \int_{x=L_{\mathrm{wall}}}^{x=L_{\mathrm{fluid}}} W^4\, dx \tag{4}$$

Next, we defined the length utilization $a_{\mathrm{L}}$ expressing the integral term from equations (3),(4) with $L$ being the total length of cantilever:

$$a_{\mathrm{L}}(L_{\mathrm{fluid}}, L) = \frac{1}{L} \int_{x=0}^{x=L_{\mathrm{fluid}}} W^4\, dx \tag{5}$$

The term $a_{\mathrm{L}}$ from equation (5) expresses the length of the cantilever that is utilized for producing $\Delta f_{rms}$, relative to the total length of $L$ the cantilever that would produce maximum $\Delta f_{rms}$ if the cantilever had uniform displacement along its length. With the term 'uniform displacement' we refer to the hypothetical case where the entire cantilever would vibrate up and down with same displacement along its length ($W(x) = 1$ for $0 \le x \le L$). The length utilization $a_L$ depends on the length of fluid channel with respect to the total length of cantilever ($L_{\mathrm{fluid}}/L$), as well as the mode number $n_{\mathrm{m}}$. Since for our designs the fluid channel extends along most of the length of the cantilever (Supporting Info S4, $L_{\mathrm{fluid}}/L > 0.95$), $a_L$ lies in the range $a_{\mathrm{L}} = 7 - 15\%$ (Supporting Info S3).

Substituting $a_{\mathrm{L}}$ from equation (5) into equations (3), (4), we expressed the two terms as:

$$\iiint_{V_{\mathrm{main}}} W^4(x) dV = 2\, H\, W\, L\, a_{\mathrm{L}}(L_{\mathrm{fluid}}, L) \tag{6}$$

$$\iiint_{V_{\mathrm{tip}}} W^4(x) dV = H\, W_{\mathrm{int}}\, L\, [\, a_{\mathrm{L}}(L_{\mathrm{fluid}}, L) - a_{\mathrm{L}}(L_{\mathrm{wall}}, L)] \tag{7}$$

Thus, we expressed the integral in equation (1) using equations (6), (7):

$$\iiint_V W^4(x) dV = H\, L\, (2\, W\, a_{\mathrm{L}}(L_{\mathrm{fluid}}, L) + W_{\mathrm{int}}\, [\, a_{\mathrm{L}}(L_{\mathrm{fluid}}, L) - a_L(L_{\mathrm{wall}}, L)]\,) \tag{8}$$

Next, to express equation (7) using a single term, we defined the volume utilization factor $a_{\mathrm{V}}$ as:

$$a_{\mathrm{V}} = \frac{H\,L\,(\;2W\,a_{\mathrm{L}}(L_{\mathrm{fluid}}, L)\;+\;W_{\mathrm{int}}[\,a_{\mathrm{L}}(L_{\mathrm{fluid}}, L) - \,a_{\mathrm{L}}(L_{\mathrm{wall}}, L)]\,)}{V} \tag{9}$$

Analogously to the term $a_{\mathrm{L}}$, the volume utilization factor $a_{\mathrm{V}}$ from equation (8) expresses the fluid volume of the cantilever that is utilized for producing $\Delta f_{rms}$, relative to the total fluid volume of $V$ the cantilever that would produce maximum $\Delta f_{rms}$ if the cantilever had uniform displacement along its length.

Thus, by inserting equation (9) into equation (1), we expressed the $\Delta f_{rms}$ as:

$$\Delta f_{rms} = \frac{f^2 c\, m_{\mathrm{AAV}}^2\, a_{\mathrm{V}} V}{4 m_{\mathrm{eff}}^2} \tag{10}$$

Next, we grouped all the variables in equation (10) related to the cantilever to define the cantilever's sensitivity $S$ as:

$$S = f^2 a_{\mathrm{V}} V / (4 m_{\mathrm{eff}}^2) \tag{11}$$

Since $f \sim 1/L$, $V \sim L^3$, $m_{\mathrm{eff}} \sim L^3$, the cantilever sensitivity exhibits the following length scaling:

$$S \sim 1/L^5 \tag{12}$$

Using equation (11), we expressed equation (10) as:

$$\Delta f_{rms} = S\,c\,m_{\mathrm{AAV}}^2 \tag{13}$$

Among all the available cantilevers, we found that the smallest one (Supporting Info S4, 0.7 × 0.7) has the highest sensitivity (Figure 1b). Using equation (8), we found that this device has a volume factor of $a_{\mathrm{V}} = 13.3\%$ which we used in equation (9) for all the experiments and simulations of the study.

To calculate AAV mass using equation (13), we use the concentration value $c$ provided either by the nominal value (Figures 1d, 2) determined by the vendor of samples, or by the measured value (Figures 3d, 4) determined by the workflow of SECMALS (Supporting Info S12).

### S3. Length utilization factor $a_\mathrm{L}$

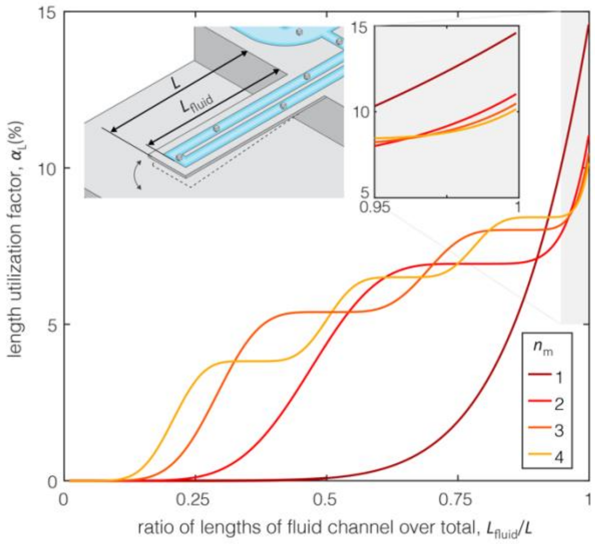


**Figure S3**: **Length utilization factor $a_\mathrm{L}$.** The term $a_\mathrm{L}$ from equation (5) (Supplementary Info S2) expresses the length of the cantilever that is utilized for producing $\Delta f_{rms}$, relative to the total length of $L$ the cantilever that would produce maximum $\Delta f_{rms}$ if the cantilever had uniform displacement along its length. The factor $a_\mathrm{L}$ depends on the length of the fluid channel $L_\mathrm{fluid}$ with respect to the total length $L$ of the cantilever (schematic in inset) and the resonant mode number $n_\mathrm{m}$. The inset plot refers to the area with the gray shading in the main plot, having axes with the same variables.

### S4. Properties of available suspended micro- and nanochannel resonator devices for measurement of AAV

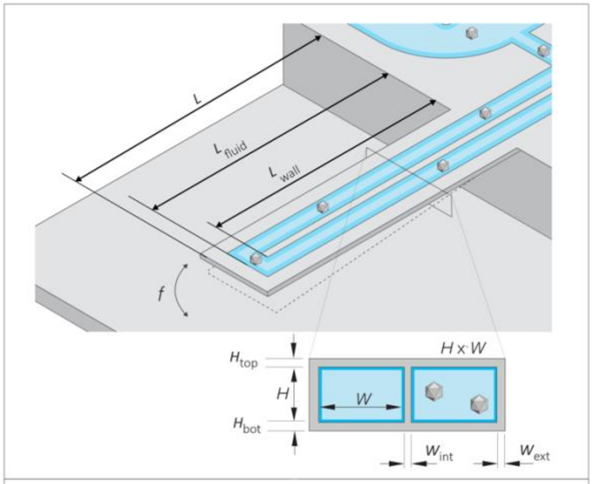


| dimensions (μm) | | | | | |
|---|---|---|---|---|---|
| name ($H$x$W$) | 0.7x0.7 | 0.7x1 | 3x3 | 8x8 | 15x20 |
| $H$ | 0.7 | 0.7 | 3 | 8 | 15 |
| $W$ | 0.7 | 1.0 | 3 | 8 | 20 |
| $H_{bot}$ | 0.2 | 0.2 | 1 | 2 | 2 |
| $H_{top}$ | 0.2 | 0.2 | 1 | 2 | 2 |
| $W_{ext}$ | 1.0 | 1.0 | 2 | 6 | 6 |
| $W_{int}$ | 0.5 | 0.5 | 1.5 | 5 | 5 |
| $L_{wall}$ | 16.3 | 31.0 | 98 | 196 | 329 |
| $L_{fluid}$ | 17.0 | 32.0 | 99 | 202 | 349 |
| $L$ | 17.5 | 32.5 | 100 | 210 | 355 |
| effective mass $m_{eff}$ (pg) & frequency $f$ (MHz) at first resonant mode | | | | | |
| $m_{eff}$ | 50 | 100 | 2,100 | 52,000 | 171,000 |
| $f$ | 4.50 | 1.20 | 0.50 | 0.38 | 0.19 |

**Figure S4 : Suspended micro- and nanochannel resonator properties**. The effective mass $m_{\mathrm{eff}}$ and resonant frequency $f$ of the cantilevers measured[21] when filled with water.

### S5. Methods

**Fabrication and design of devices.** The suspended nanochannel resonator devices (SNR) were fabricated at CEA Leti (Grenoble, France) using 8-inch. (200mm) silicon wafer technology. The technology enables the cantilever of each device to oscillate in a dedicated vacuum cavity containing an on-chip getter to maintain the high vacuum, thus ensuring high quality factor during operation. The type of devices used in this study (Supporting Info S4, $0.7 \times 0.7$), has one cantilever with four fluidic ports drilled on the top glass wafer to access two bypass channels connected to the inlet and the outlet of each cantilever.

**Operation of devices.** Each SNR device is actuated at the first vibration mode by a piezo-ceramic plate on top of which the device is epoxy-bonded. A dedicated phase-locked loop (PLL) in closed loop maintains the cantilever at resonance. Precision pressure regulators (electronically controlled Proportion Air QPV1 and manually controlled Omega PRG101-25) were used to flow particle solutions within each device. To measure the signal of change in resonance frequency, $\Delta f$, optical readout method was employed with a field programmable gate array (FPGA, Altera Cyclone IV on DE2-115) connected via ethernet cable to a desktop computer. The experiments were performed using a custom code written in LabVIEW 2017 software.

**Tested solutions of gold calibration nanoparticles.** Solutions of gold nanoparticles were purchased from BBI Solutions (Crumlin, UK) for two different nominal diameters: i) $d_{\text{nom}} = 5$ nm (EM.GC10/4, batch #19100149) and concentration $c = 50 \times 10^{12}$ particles/mL and ii) $d_{\text{nom}} = 20$ nm (EM.GC20, batch #15022) and concentration $c = 0.7 \times 10^{12}$ particles/mL.

**Tested solutions of viral vectors.** Solutions of AAV of serotype 5 (AAV5) were purchased from Virovek (Hayward, CA USA) including AAV5-GFP (Lot: 17-598) with genetic construct of green fluorescent protein (GFP) corresponding to nominal DNA bases $n_{\text{DNA}} = 3.3$ kilobases, AAV5-empty (Lot: 19-528E) with $n_{\text{DNA}} = 0$ kilobases and AAV5-4891 (Lot: 19-152) with $n_{\text{DNA}} = 4.9$ kilobases at nominal concentrations of $c_{nom} = 20 \times 10^{12}$ particles/ml.

**Preparation of solution**: AAV and gold nanoparticle solutions were diluted using PBS pH 7.4 (1X) (Gibco, Ref: 10010-023, Lot: 2198731) filtered with Sterile Syringe Filter 0.2 µm Cellulose Acetate (VWR, PN: 28145-477. LOT: FE 3644). Dilutions were done at 2X, 4X,8X, 10X, 20X.

**Cleaning of SNR devices**: Devices were occasionally clogged due to their small cross-section area ($700\ x\ 700\ nm^2$) of the fluidic nanochannel. To unclog the nanochannel, aqueous solution of 5% w/v Tergazyme was used interchangeably with aqueous solution of 10% v/v Bleach, as well as Isopropanol (IPA) and Acetone.

**Analytical Ultracentrifugation (AUC):** Absorbance of samples was measured using a NanoDrop. All samples were diluted to 0.6 Au (Absorbance units) at 280 nm. 400 µL of 1X PSB and pluronic F68 and 400 µL of sample were loaded into two-sector velocity cells. Loaded cells were placed in an 8-hole rotor. The filled rotor was placed in the AUC Optima model and temperature-equilibrated at 20°C for at least 2 hours. Following temperature-equilibration, radial scans at 15,000 rpm were initiated with 150 scans per sample. Data were collected using three signals; 280 nm, 260 nm and RI. Data were analyzed in the program Sedfit using the c(s) model. Results for weighted values were extracted using data from 280 nm and RI.

**Analytical Anion Exchange (AEX):** AEX was performed on an Agilent 1260 Infinity II LC System with a Sepax Proteomix WAX-NP5 (weak anion exchange) chromatography column. The system was equilibrated with 50 mM sodium acetate, 20 mM Bis tris propane (BTP), pH 8.0 and 20 µL of each sample was injected onto the column. The column was washed with the above buffer for 10 minutes at 0.5 mL/min and samples were eluted by a steady gradient to 0.25 M sodium acetate, 20 mM BTP, pH 8.0 at 25 minutes followed by a steady gradient to 1 M sodium acetate, 20 mM BTP, pH 8.0 at 45 minutes. UV absorbance (214, 230, 260, 280 nm) and fluorescence (Ex = 280 nm, Em = 340 nm) were monitored. Chromatograms were exported and curve fitting was performed in OriginPro 2021b. The complete methodology is presented in Supporting Info S14.

**Size Exclusion Chromatography Multi-Angle Light Scattering (SECMALS)**: Analysis was performed according to previous methodology.[7] A Sepax SRT SEC-1000 column with guard column was used for all experiments. The column was equilibrated with an isocratic elution buffer of PBS (2X) + 10% EtOH for at least 12 hours. 20 µL of sample was injected for each analysis. UV absorbance of column eluates at 260 nm and 280 nm was detected by a multiple-wavelength diode array detector, light scattering was detected by a Wyatt HELIOS, and refractive index detected by Wyatt TREX. Data analysis was done on Wyatt ASTRA 8 software.

**Dynamic Light Scattering (DLS)**: DLS measurements were performed on a Wyatt Dynapro plate reader III. For each measurement, 30 µL of neat sample was pipetted into one well of a 384-well plate (Aurora Microplates, ABM210100). A total of 30 acquisitions lasting 1 second each were taken per well. To remove readings contaminated by dust or other very large particulates, a data filter was applied to exclude acquisitions with baseline >0.005 arbitrary units of autocorrelation intensity. The remaining acquisitions were averaged to give results for each sample.

**Alkaline Agarose Gel Electrophoresis (AAGE):** Encapsulated genome size distribution was confirmed by 1% agarose gel electrophoresis, staining with SYBR Gold (SG) Nucleic Acid Gel Stain (Invitrogen), and visualizing under UV-light using a Protein Simple FluorChem Imager. Agarose gel was prepared by dissolving 2 g SeaKem LE agarose in 200 mL 1x alkaline buffer (50 mM NaOH, 1 mM EDTA). Before electrophoresis, $1 \times 10^{11}$ vg of samples were added to 1.5 μL 10% SDS (Ambion) and 7.5 μL 6x Alkaline Gel Loading Dye (Alfa Aesar) and brought to 25 μL with 1x alkaline buffer. 7.5 μL of 1 kb ladder (New England Biolabs) was added to 6 μL 10% SDS, 30 μL 6x alkaline loading dye, and 31.5 μL of 1x alkaline buffer for triplicate runs. 18 μL of samples and ladders were loaded on the gel, which was resolved using 65 V/cm for 4 hours, before three 10-minute washes in neutralizing buffer (1 M Tris-HCl, 1 M NaCl, pH 7.40). DNA on the gel was stained with SG solution (30 μL SG, 20 mL neutralizing buffer, 180 mL Milli-Q water for 1 hour, followed by a 5 and then 55 minute wash in neutralizing buffer. DNA was visualized on a Protein Simple FluorChem Imager using the Ethidium Bromide setting.

**Post processing of experimental data from the SNR.** Data from the SNR were processed using MATLAB 2019b with the Signal Processing Toolbox™.

### S6. Dynamic Light Scattering (DLS) for characterizing size of gold nanoparticles, and aggregation of AAV

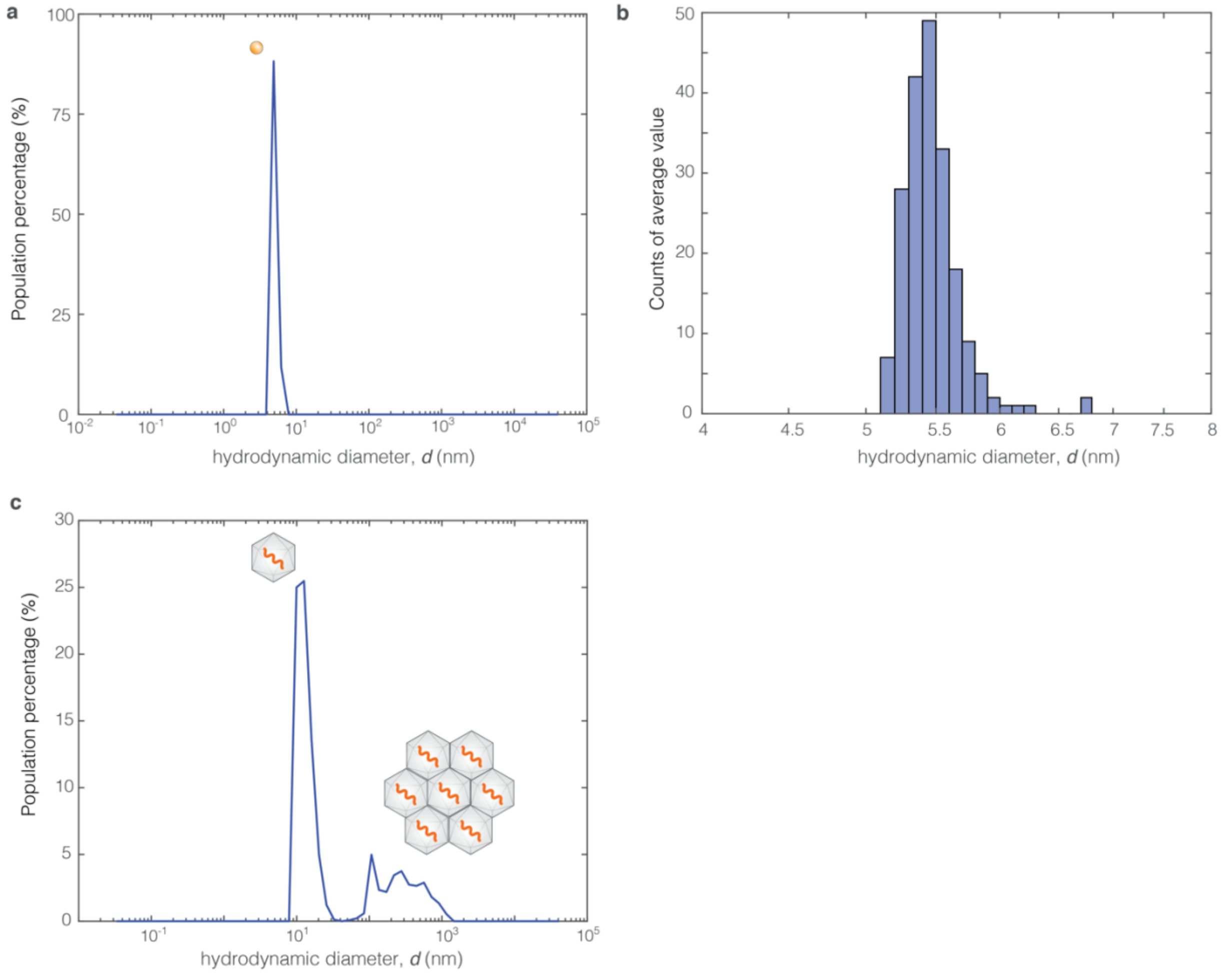


**Figure S6| Dynamic light scattering of nanoparticles. a,** Example curve of population percentage of calibration gold nanoparticles (inset is schematic of gold nanoparticle) of nominal diameter $d_{\mathrm{Au,nom}} = 5$ nm. The solution of calibration gold nanoparticles was found monodisperse. **b**, Histogram of mean values extracted from curves such as the curve of the population percentage in panel **a** ($n = 198$ curves) yielding a mean hydrodynamic diameter of $d = 5.4$ nm with standard deviation $\sigma_d = 0.2$ nm. The mass of gold nanoparticles was calculated as $m_{\mathrm{Au,ref}} = (\rho_{Au} - \rho)\pi d^3/6$ and was used as a reference for comparison with the measured mass from nanochannel resonator (Figure 2). The calculation was done using the values $\rho_{Au} = 19{,}320\ \mathrm{kg/m^3}$ for gold nanoparticles and $\rho = 1{,}000\ \mathrm{kg/m^3}$ for aqueous solution. **c**, Curve of population percentage of AAV5-GFP indicates a peak corresponding to the nominal diameter of AAV ($d = 20$ nm) indicating single AAV capsid, as well as a population of aggregates ($d > 100$ nm). The schematics of single AAV and aggregates are shown as insets.

### S7. Computational model for estimating the accuracy and precision of experimental measurement

In addition to theory (Supporting Info S2), we developed a computational model to gain physical intuition about our experimental measurements, and computationally estimate its accuracy and precision.

The flow of particles, either AAV or gold nanoparticles, is characterized by the interplay of advection (transport of particles as they flow through the cantilever due to pressure-driven flow) and diffusion (three-dimensional Brownian motion under confinement). The flow in the cantilever is a laminar, low Reynolds number flow (see table in this section) with a three-dimensional parabolic profile of flow speed[25]. Given an average flow speed $V_{\mathrm{flow}}$, we assumed an advection time scale, or transit time $\tau_{\mathrm{a}} = 0.05 - 1.0$ s which refers to the average time is takes a given nanoparticle to travel through the cantilever. The diffusion time scale is set by the diffusivity coefficient $D = k_{\mathrm{B}}T/(3\pi\mu d)$, where $k_B = 1.380649 \times 10^{-23}$ J/K is Boltzman constant, $T$ is temperature of fluid, $\mu$ is dynamic viscosity of fluid, and $d$ is diameter of nanoparticles. Thus, for AAV the diffusion time scale is $\tau_d = W^2/(6D) = 0.0037$ s, in which $W$ is the characteristic dimension of the square cross-section of the flow channel (Supporting Info S4), and the numerical prefactor accounts for the three-dimensional diffusion. The ratio of the two time scales result in a Péclet number of $Pe = \tau_{\mathrm{a}}/\tau_{\mathrm{d}} = 0.0037 - 0.37$ (Table of this section**Error! Reference source not found.Error! Reference source not found.**) indicating that the effect of diffusion is dominant over advection along the length of the fluid channel.

Thus, our model captures the interplay of both advection and diffusion of AAV particles within the channel. We deterministically modeled the advection of AAVs in a pressure-driven flow as a laminar, low Reynolds flow characterized by a 3-D parabolic velocity profile.[25] We modeled Brownian motion of the AAVs due to diffusion as a random walk characterized by the diffusion coefficient $D$. In the interest of simplicity, we assumed there is no adhesion to the walls or interaction between particles; we only considered the elastic collision of the particles on the walls of the channel.

Taking all of the above, we modelled the entry of particles in the inlet of the fluid channel as a Poisson distribution:

$$P(k) = \frac{\lambda\, dt\, e^{-\lambda dt}}{k!} \tag{14}$$

Equation (14) defines the probability that $k$ particles enter the cantilever after the elapse time of $dt$, given an entry rate $\lambda = cHWV_{flow}$ of particles entering per unit time, where $c$ is concentration, and H, W are the dimensions of the cross-section area of the fluid channel (Supporting Info S4). As the particles travel through the channel advecting and diffusing, they exit the cantilever with no locally imposed condition at the outlet. Applying this approach of stochastic modelling, we simulated the flow of AAVs through the cantilever resulting at any given time $t$ in a number $n_{\mathrm{p}}(t)$ of particles present in the channel at three-dimensional positions $\bar{r}_{\mathrm{i}}(t) = \big(x_{\mathrm{i}}(t), y_{\mathrm{i}}(t), z_{\mathrm{i}}(t)\big)$ where i is the index of the particles $\mathrm{i} = 1,2,\ldots,n_{\mathrm{p}}(t)$.

Therefore, by recording the trajectory $\bar{r}_{\mathrm{i}}(t)$ of each AAV through the channel, we derived the resonant frequency change contributed by each of the $n(t)$ AAV present in the fluid channel. In doing so, we calculated the net resonant frequency signal $\Delta f(t)$ as the sum of these individual contributions, scaled by device characteristics (Supporting Info S4, $m_{\mathrm{eff}}, f$), with the optional addition of noise $\Delta f_{\mathrm{noise}}(t)$:

$$\Delta f(t) = -\frac{f}{2m_{\mathrm{eff}}} \sum_{i=1}^{n_{\mathrm{p}}(t)} W^2(x_{\mathrm{i}}(t))\, m_{\mathrm{AAV}} + \Delta f_{\mathrm{noise}}(t) \tag{25}$$

Notably, in equation (15), only the longitudinal position ($x_{\mathrm{i}}(t)$ i.e. along length $L$, Supporting Info S4) of the particle matters for the generated signal $\Delta f(t)$. We modelled the noise term $\Delta f_{\mathrm{noise}}(t)$ as colored noise of the form $\Delta f_{\mathrm{noise}}(\xi) = \beta/\xi^{\alpha}$ based on experimental study of noise of our system that showed that colored noise was a good approximation (Supporting Info S9). Furthermore, we set the $rms$ of $\Delta f_{\mathrm{noise}}$ according to the experimental measurements of noise (Supporting Info S9).

Lastly, we set the solution of particles to either have a uniform mass distribution such as gold nanoparticles of $d = 5.4$ nm and $m_{Au} = 1.51$ ag (Figure 2) or a non-uniform mass distribution such as in AAV solution (Figure 3c) as determined by Analytical Ultracentrifugation (Figure 3b). All the parameters used as inputs in the model are summarized in the table of this section.

| Parameters used as inputs in computational model | | | | | | |
|---|---|---|---|---|---|---|
| Parameter | Symbol | Type | Units | Minimum value (if range) | Maximum value (if applicable) | Notes |
| density of media | $\rho$ | Cantilever geometry, fluid properties and simulation parameters | $\mathrm{kg/m^3}$ | 1000 | | Value used for aqueous solution, in particular PBS (Phosphate buffer solution) |
| dynamic viscosity of media | $\mu$ | | Pa s | $8.9 \times 10^{-4}$ | | For same fluid as in ρ |
| temperature | $T$ | | K | 298 | | Room temperature |
| cross-sectional area of nanochannel | $H \times W$ | | $\mu\mathrm{m}^2$ | $0.7 \cdot 0.7$ | | Corresponds to cantilever device $0.7 \times 0.7$ (Supporting info S4) |
| average length of transit path through Nanochannel | $L_{\mathrm{transit}} = L_{\mathrm{fluid}} + L_{\mathrm{wall}} + W + W_{\mathrm{int}}$ | | µm | 33.3 | | Corresponds to cantilever device $0.7 \times 0.7$ (Supporting info S4) |
| transit time or advection time scale | $\tau_{\mathrm{a}}$ | | s | 0.01 | 1.0 | |
| average fluid velocity | $V_{\mathrm{flow}} = L_{\mathrm{transit}}/\tau_{\mathrm{a}}$ | | µm/s | 33.3 | 660 | |
| time step | dt | | s | 0.001 | | Used for all simulations |
| entry rate $\lambda$: Number of particles entering cantilever per unit of time | $\lambda = c\ H\ W\ V_{\mathrm{flow}}$ | | particles /$s$ | 16.3 | 6468 | Entry rate used in equation (14) |
| Reynolds number | $\mathrm{Re} = V_{\mathrm{flow}} W/\mu$ | | - | $2.7 \times 10^{-5}$ | $5.4 \times 10^{-4}$ | |
| average concentration | $c$ | Solution of Adeno-Associated Viruses (AAV) | particles /ml | $1 \times 10^{12}$ | $2 \times 10^{13}$ | |
| buoyant mass of AAV | $m_{\mathrm{AAV}}$ | | ag | 1.26 | 2.33 | Limit of range defined from mass conversions (Supporting Info S1). Distribution determined by Analytical Ultracentrifugation (Figure 3b) |
| diameter of AAV | $d$ | | nm | 22 | | |
| diffusivity coefficient of AAV | $D = \frac{k_{\mathrm{B}}T}{3\pi\mu d}$ | | $\mathrm{m^2 s}$ | $2.23 \times 10^{-11}$ | | |
| diffusion time scale of AAV | $\tau_{\mathrm{d}} = W^2/(6D)$ | | s | 0.0037 | | |
| Péclet number of AAV | $\mathrm{Pe} = \tau_{\mathrm{d}}/\tau_{\mathrm{a}}$ | | - | 0.0037 | 0.37 | |
| average concentration | $c$ | Solution of gold nanoparticles (Au) | particles /ml | $1 \times 10^{12}$ | $2 \times 10^{13}$ | |
| buoyant mass of Au | $m_{\mathrm{Au}}$ | | ag | 1.5 | | Simulation run (Figure 2) with same $m_{\mathrm{Au}}$ for all particles |
| diameter of Au | $d$ | | nm | 5.4 | | |
| diffusivity coefficient of Au | $D = \frac{k_{\mathrm{B}}T}{3\pi\mu d}$ | | $\mathrm{m^2 s}$ | $9.08 \times 10^{-11}$ | | |
| diffusion time scale of Au | $\tau_{\mathrm{d}} = W^2/(6D)$ | | s | 0.0009 | | |
| Péclet number of Au | $\mathrm{Pe} = \tau_{\mathrm{d}}/\tau_{\mathrm{a}}$ | | - | 0.0009 | 0.09 | |

**Table | Parameters of computational model.** This table contains all the values used for simulation juxtaposed with experiments in Figures 2,3,4.

### S8. Frequency domain study of simulated signal $\Delta f(t)$ due to nanoparticles in the absence of noise

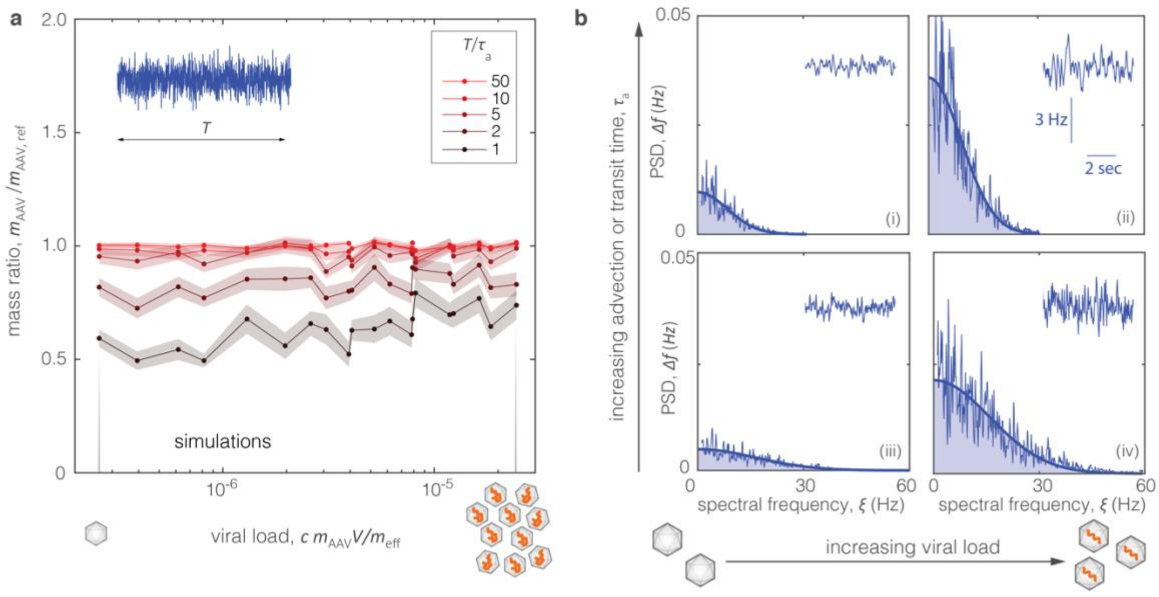


**Figure S8 | Frequency domain study of simulated signal exclusively due to AAV. a,** Mass ratio of AAV mass $m_{\text{AAV}}$ extracted from $\Delta f_{rms}$ of simulated signals over $m_{\text{AAV,ref}}$ vs the viral load $c\ m_{\text{AAV}}V/m_{\text{eff}}$ . The viral load represents the total mass of viruses in cantilever, normalized by the effective mass of the cantilever. Transit time is $\tau_{\text{a}} = 0.1$ s. The range of viral load spans from empty or 'light' capsids ($m_{\text{AAV}} = 1.26$ ag) at concentrations of $c = 10^{12}$ particles/ml to full or 'heavy' capsids ($m_{\text{AAV}} = 2.31$ ag) of concentrations at $c = 20 \times 10^{12}$ particles/ml (Supporting Info S1), qualitatively indicated by schematics below horizontal axis. Every point is average $n = 30$ time traces of duration 30 sec each. The colored bands define standard error around each point. $T$ denotes the duration of signal used for calculating $\Delta f_{rms}$ and $\tau_{\text{a}}$ the advection or 'transit' time of a given nanoparticle through the cantilever. **b,** Power Spectral Density (PSD) for simulated signals vs the spectral frequency $\xi$. The thick blue line denotes Gaussian fit $\Delta f(\xi) = \gamma e^{-\delta \xi^2}$. The area under $\Delta f(\xi)$ is equal to $\Delta f_{rms}$ (blue shade). The area increases for increasing viral load (e.g. i to ii, or iii to iv) but stays the same for increasing transit time $\tau_{\text{a}}$, corresponding to slower flow (e.g. iii to I, or iv to ii ).

### S9. Spectral denoising method for neutralizing effect of noise when calculating nanoparticle mass

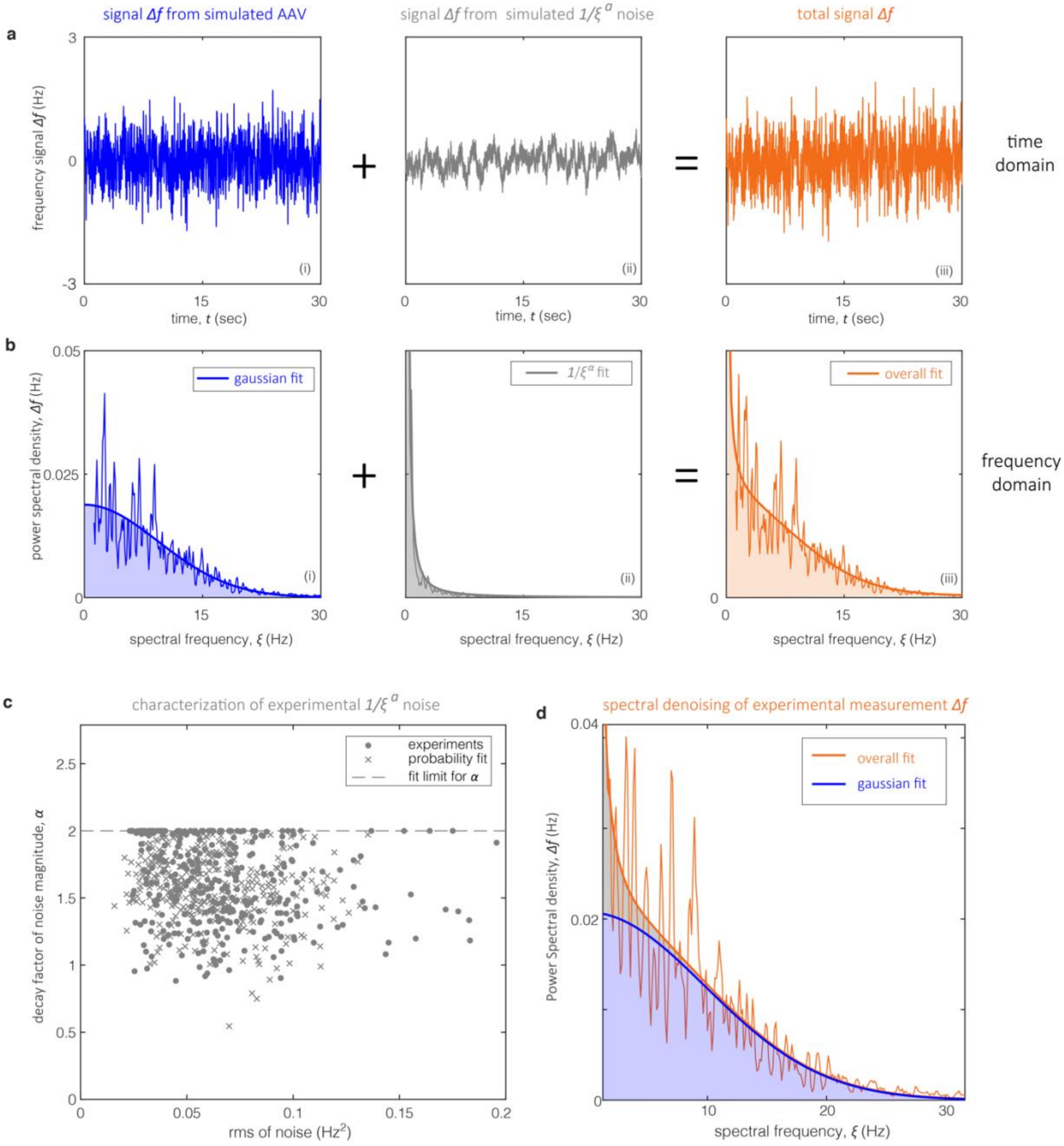


**Figure S9 | Spectral denoising of experimental data. a,b,** Simulations of signals $\Delta f$ generated by the 'pure signal' by viruses in the absence of noise (i), and by colored $1/\xi^{\alpha}$ noise (ii) in both the time (**a**) and frequency domain as P-welch Power Spectral Density (**b**). The total signal (iii) is summation of (i) and (ii). The thick curves in (**b**) denote fits based on canonical forms of Gaussian $\Delta f(\xi) = \gamma e^{-\delta\xi^2}$(i) (Supporting information S8) and $\Delta f_{noise}(\xi) = \beta/\xi^{\alpha}$ (ii) in the frequency domain. The colored areas denote areas under the fitted curves which are equal to the root mean square $rms$ of each signal. **c**, Experiments for characterization of noise by measuring $rms$ when flowing buffer solution without nanoparticles in the cantilever. Assuming colored noise, we calculated the fit $\Delta f(\xi) = \beta/\xi^{\alpha}$ of the Power Spectral Density where $\alpha = [-2\ 2]$. Crosses denote probability fits for the pairs $(a, rms)$. **d**, Spectral denoising of experimental measurement consisting of applying the fit $\Delta f(\xi)$=$\gamma e^{-\delta\xi^2} + \beta/\xi^{\alpha}$ on the Power Spectral Density of $\Delta f$, and then calculating $\Delta f_{rms}$ by accounting only for the area under the Gaussian portion of the fit as $\Delta f_{rms} = 0.5\gamma\sqrt{\pi/\delta}$. Calculating the $\Delta f_{rms}$ with spectral denoising calculates the expected mass of calibration gold nanoparticles (Figure 2).

### S10. Resolution of mass measurement

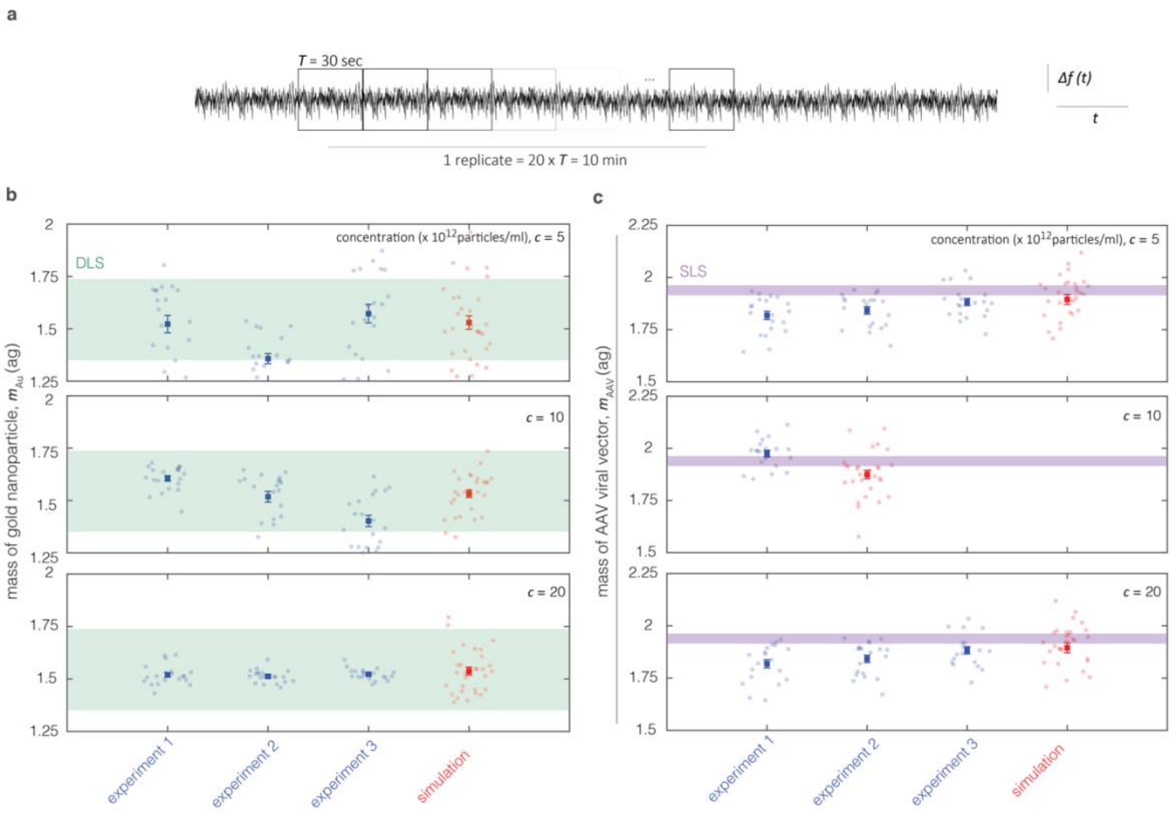


**Figure S10 | Resolution of mass measurement defined as standard error over finite time sampling window. a,** Schematic of frequency signal $\Delta f(t)$. The spectral denoising method is applied to sampling windows of $T = 30$ sec (squares). Twenty sampling windows define one replicate of 10 min . **b,** Measured mass of gold nanoparticles (Figure 2) from both experiments (exp, blue) and simulations (sim, red) where error bars denote standard error for sampling windows of 10 min. The green bands denote range of measurement from Dynamic Light Scattering (DLS). The standard error lies below 100 zg for concentrations of $c = 5 \times 10^{12}$ particles/ml, going down to near 10 zg for concentrations of $c = 20 \times 10^{12}$ particles/ml. **c,** Measured mass of AAV viral vectors (Figure 3d) with same notation as in panel **b**. The purple bands denote range of measurement from Static Light Scattering (SLS). The standard error lies near 10 zg for the three concentrations.

### S11. Analytical Ultracentrifugation (AUC) of AAV samples

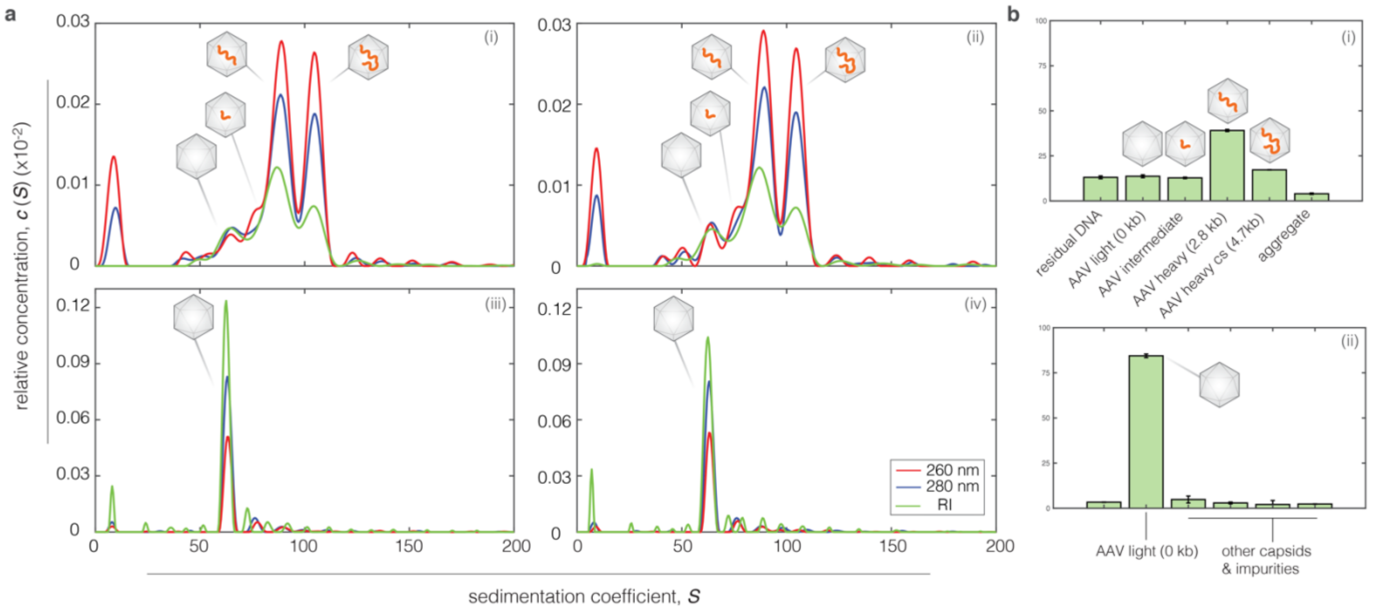


**Figure S11 | Analytical Ultracentrifugation of samples. a,** Analytical Ultracentrifugation (AUC) analysis of AAV-GFP (i,ii) and AAV-empty (iii,iv) using Refractive Index (RI) signal (green) and 260, 280 nm UV wavelengths (red, blue) (i). $c_{260}/c_{280} > 1$ indicates AAV capsids filled with DNA[26] (e.g., 'heavy capsids'). **b,** Population percentages extracted from panel **a** with numbers corresponding to peaks indicating different types of AAV (insets) for AAV-GFP (i) and AAV-empty (ii). The letters 'cs' in 'AAV heavy cs'denote complementary strand, filling up the capsid to its maximum holding capacity (4.7 kb).

### S12. Measuring concentration of AAV samples

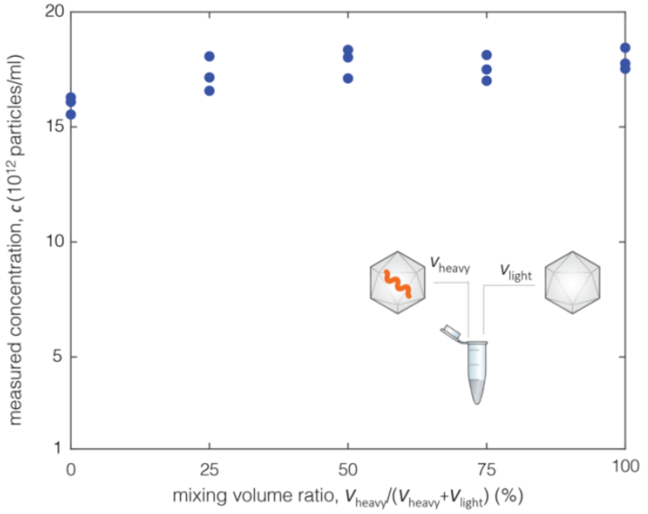


**Figure S12 | Measuring concentration of AAV samples.** Measured concentration $c$ vs. mixing volume ratio for experiments, determined by the workflow of Size Exclusion Chromatography Multi-Angle Light Scattering (SECMALS) (Supporting Info S5). For each mixing ratio, three measurements were conducted. These values were used to calculate AAV mass in Figures 3d, 4.

### S13. Limit of detection for measuring mass of AAV

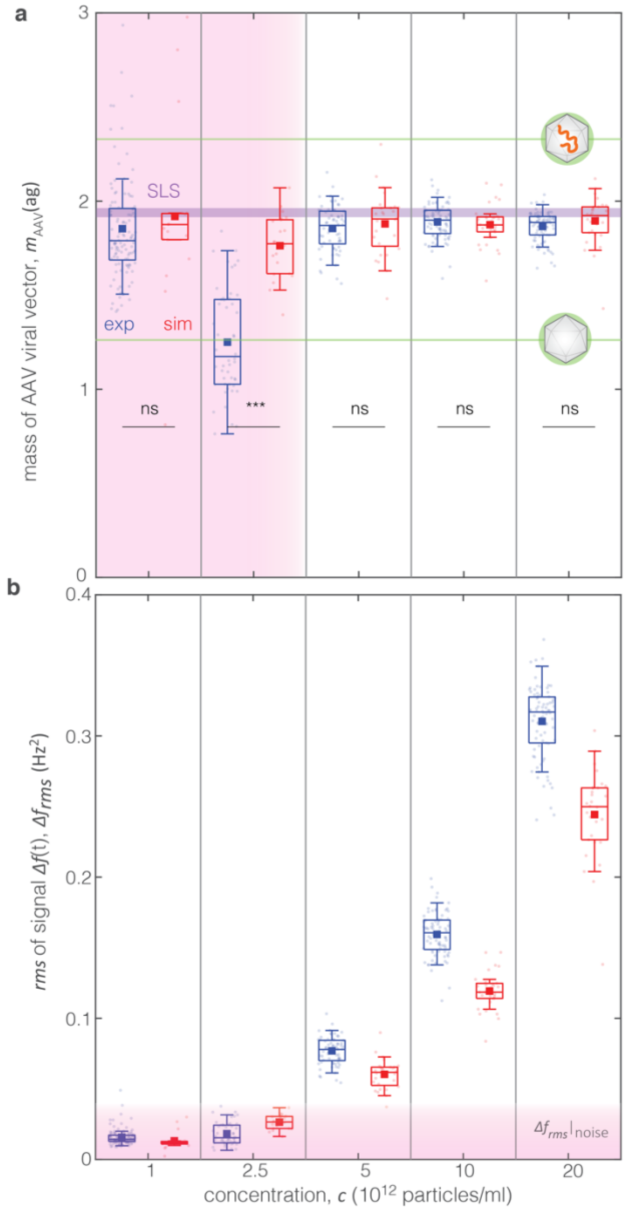


**Figure S13 | Limit of detection for measuring mass of AAV. a,** Mass $m_{\mathrm{AAV}}$ of AAV viral vector with genetic construct of GFP vs concentration $c$ (non-linear scale) specified with denoising in experiments (exp: blue) simulations (sim: red). Same data and notation as in Figure 3d but extending concentration down to $c = 1 \times 10^{12}$ particles/ml. The shaded pink area indicates the limit of detection due to the presence of significant noise, which compromises the effectiveness of the denoising technique. **b,** Measured $\Delta f_{rms}$ used to derive the mass $\mathrm{m_{AAV}}$, after applying spectral denoising (Supporting Info S9) to the signal $\Delta f(t)$ from both experiments (blue) and simulations (red). The pink shaded region denotes the $\Delta f_{rms}\,|_{\,\mathrm{noise}}$ exclusively due to noise based on the typically encountered levels of noise (Supporting Info S9). The denoising method does not work in the pink shaded region due to the noise $\Delta f_{rms}\,|_{\,\mathrm{noise}}$ being same order of magnitude as the nanoparticle $\Delta f_{rms}$, thus compromising the accuracy of measurement for concentrations below $c < 5 \times 10^{12}$ particles/ml. The difference in the $\Delta f_{rms}$ between experiments and simulations are attributed to use of different effective mass $m_{\mathrm{eff}}$ in simulations than experiments.

### S14. Anion Exchange Chromatography for characterizing AAV

This section describes the detailed methodology of analytical Anion Exchange Chromatography for characterizing AAV as opposed to a briefer description of the technique in the Methods section (Supporting Info S5).

Analytical Anion Exchange Chromatography (AEX) was performed on an Agilent Series 1260 Infinity II LC System (Agilent, Waldbronn, Germany) equipped with a binary pump, temperature-controlled 1290 series autosampler, fluorescence detector, and multiple-wavelength photodiode array detectors. The HPLC was controlled by ChemStation OpenLab LC systems software, version 2.1.1.13. 20 µl of Light and Heavy capsid sample—at $3.6 \times 10^{13}$ particles/ml and $2 \times 10^{13}$ particles/ml, respectively—was each injected in triplicates on the Proteomix WAX-NP5 4.6×250 mm column (Sepax, Newark, DE) pre-equilibrated with 20mM Bis-Tris Propane (BTP) pH 8.0, 50 mM sodium acetate (EQ buffer) for 30 min at a 0.5ml/min flowrate. The column was then washed with EQ buffer for 10min followed by a 30-minute AAV capsid elution step using a 50-250 mM linear gradient of sodium acetate, 20 mM BTP pH 8.0. Next, column-bound sample aggregates were washed off with 1 M sodium acetate, 20 mM Bis-Tris Propane (BTP) pH 8.0 (20 min; strip wash) and the column was equilibrated with the EQ buffer for 30 min prior to subsequent sample injection. Total method runtime was 90 min, including sample injection, column wash with EQ buffer, elution, strip wash, and column equilibration.

Capsid elution profiles were monitored by UV absorbance at 214, 230, 260, 280 nm and intrinsic protein fluorescence ($\lambda_{EM} =$ 280 nm and $\lambda_{EM} = 340$ nm) often used to quantify AAV capsid populations using chromatography-based methods[4]. All chromatograms obtained using fluorescence detection were analyzed using OriginPro software (Version 2020b, OriginLab Corporation, Northampton, MA, USA.). After baseline subtraction, the second derivative of each chromatogram was filtered using Savitzky-Golay algorithm and then used to deconvolute capsid elution profiles into individual peaks: a single peak for the light capsids and up to seven additional peaks for the remaining portion of the chromatograms. Iterative curve fitting was performed using a Levenberg-Marquardt (L-M) algorithm for nonlinear least-squares minimization until model convergence with a $\chi^2$ tolerance value of $10^{-6}$. Reduced $\chi^2$ is obtained by dividing the residual sum of squares (RSS) by the degrees of freedom (DOF).

For light capsid elution profiles, the main peak was globally analyzed using an exponentially modified Gaussian function (GaussMod)— widely used for asymmetric peak approximation in chromatography[26] —given by:

$$f(x) = y_0 + (f_1 * f_2)(x) \tag{16}$$

where:

$$f_1(x) = \frac{A}{t_0} e^{-\frac{x}{t_0}} \tag{17}$$

$$f_2(x) = \frac{1}{\sqrt{2\pi}\, w} e^{-\frac{(x-x_c)^2}{t_0}} \tag{18}$$

$$z = \frac{x - x_c}{w} - \frac{w}{t_0} \tag{19}$$

In equations (17)–(19), the $y_0$ denotes baseline, $A$ area, $x_c$ center, $w$ width, and $t_0$ unknown.

The obtained light capsid peak parameters were used within experimentally determined bounds for the peak center and width to model the light capsid peak in the heavy capsid elution profiles. Up to seven Gaussian peaks without parameter constraints were used to approximate heavy capsids and the strip peak, resulting in several models with a total of 4–8 peaks (see table of this section). The percentage of light capsids was estimated for each model by dividing the peak area for the light capsids by the total peak area. To probe the effects of peak fitting parameters on percentage of light capsids for the heavy capsid chromatograms, the model with five peaks was further constrained using arbitrary bounds for the Gaussian peak center ($x_c$) and width ($w$). Overall, best-fit models revealed that the heavy capsid sample contains ~10–14% light capsids, consistent with the results obtained by analytical ultracentrifugation.

| | Heavy Capsid sample | | | | | | | | | |
|---|---|---|---|---|---|---|---|---|---|---|
| | Run 1 | | | Run 2 | | | Run 3 | | | Runs 1–3 |
| Total no. of fitted peaks<br>Heavy/strip peak constraints | Reduced $\chi^2$ | Adjusted $R^2$ | Light Capsids (%) | Reduced $\chi^2$ | Adjusted $R^2$ | Light Capsids (%) | Reduced $\chi^2$ | Adjusted $R^2$ | Light Capsids (%) | Light Capsids (%) |
| 4<br>None | 0.056 | 0.994 | 7.9 | 0.101 | 0.989 | 10.8 | 0.106 | 0.988 | 11.1 | 9.9 ± 1.4 |
| 5<br>None | 0.039 | 0.996 | 10.1 | 0.041 | 0.995 | 10.2 | 0.044 | 0.995 | 10.6 | 10.3 ± 0.2 |
| 6<br>None | 0.036 | 0.996 | 10.0 | 0.038 | 0.996 | 10.7 | 0.042 | 0.995 | 11.2 | 10.6 ± 0.6 |
| 7<br>None | 0.040 | 0.996 | 11.4 | 0.036 | 0.996 | 10.6 | 0.040 | 0.996 | 11.1 | 11.0 ± 0.3 |
| 8<br>None | Fit did not converge; $\chi^2$ tolerance value of $10^{-6}$was not reached | | | | | | | | | |
| 5<br>$x_c$ ± 0.2min | 0.052 | 0.994 | 12.062 | 0.053 | 0.994 | 12.0 | 0.060 | 0.993 | 12.8 | 12.4 ± 0.4 |
| 5<br>$x_c$ ± 0.1min | 0.058 | 0.994 | 11.442 | 0.060 | 0.993 | 11.9 | 0.065 | 0.993 | 12.2 | 11.9 ± 0.3 |
| 5<br>$x_c$ ± 0.05min | 0.077 | 0.992 | 11.796 | 0.081 | 0.991 | 12.3 | 0.069 | 0.992 | 12.1 | 12.1 ± 0.2 |
| 5<br>$x_c$ ± 0.05min,<br>w<5min | 0.075 | 0.992 | 12.3 | 0.080 | 0.991 | 12.9 | 0.071 | 0.992 | 12.8 | 12.6 ± 0.3 |
| 5<br>$x_c$ ± 0.05min,<br>w<4min | 0.077 | 0.991 | 12.7 | 0.081 | 0.991 | 13.3 | 0.074 | 0.992 | 13.0 | 13.0 ± 0.2 |
| 5<br>$x_c$ ± 0.05min,<br>w<3min | 0.081 | 0.991 | 12.5 | 0.083 | 0.991 | 13.3 | 0.080 | 0.991 | 13.2 | 13.0 ± 0.3 |
| 5<br>$x_c$ ± 0.05min,<br>w<2min | 0.073 | 0.992 | 13.6 | 0.072 | 0.992 | 14.1 | 0.077 | 0.992 | 14.5 | 14.1 ± 0.4 |
| 5<br>$x_c$ ± 0.05min,<br>w<1min | 0.358 | 0.960 | 17.6 | 0.356 | 0.960 | 18.1 | 0.379 | 0.959 | 18.7 | 18.1 ± 0.4 |

**Table | Data from Anion Exchange Chromatography**. Adjusted $R^2$ is a modified version of $R^2$, adjusted for the number of predictors in the fitted line.

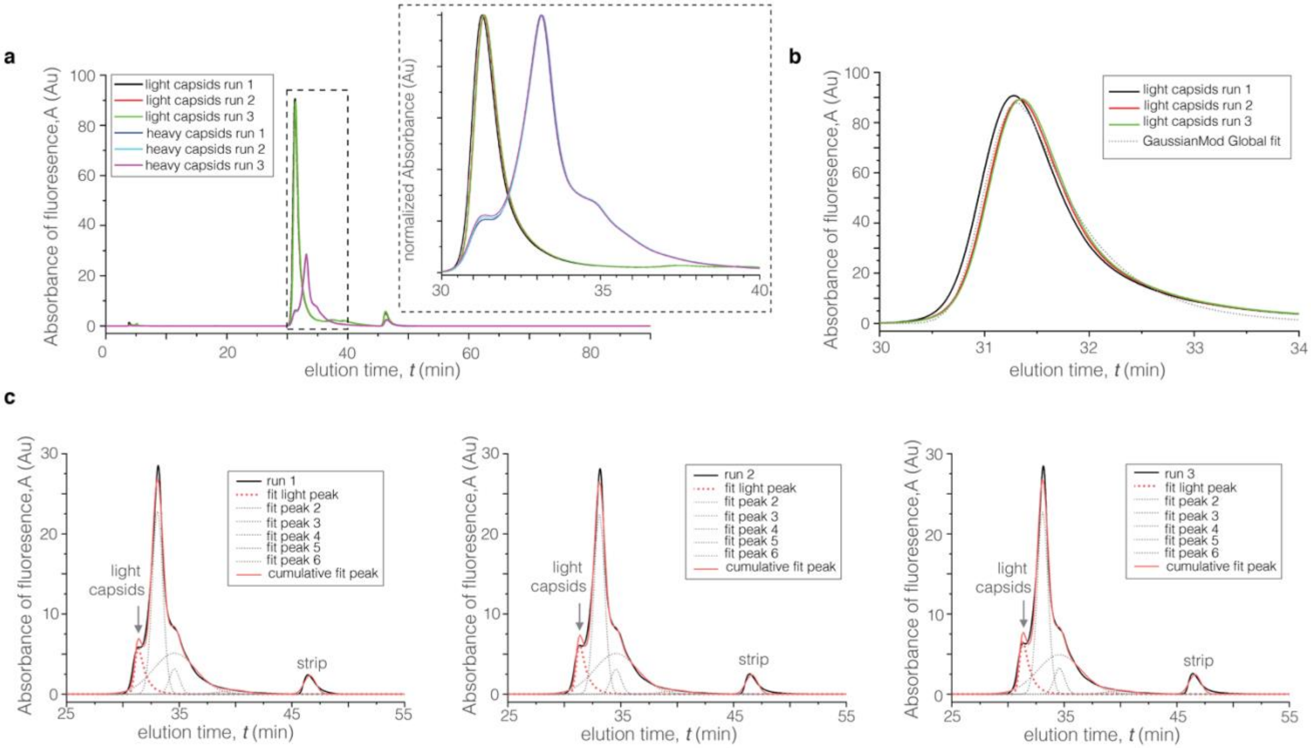


**Figure S14 | Visualization of vector genome distribution by analytical anion exchange chromatography (AEX)**. **a,** Comparison of light and heavy AAV5 capsid chromatograms using intrinsic protein fluorescence detection with $\lambda_{EM} = 280$ nm and $\lambda_{EM} = 340$ nm. Capsid elution times correlate with the amount of encapsidated DNA, with lighter capsids eluting prior to heavier capsids containing higher amount of DNA. **b,** Global analysis of empty AAV5 capsid chromatograms using an exponentially modified Gaussian peak function (GaussMod). **c,** Analysis of heavy AAV5 capsid elution profiles using a non-linear peak-fitting algorithm, shown for three replicate injections (left, middle, and right panels). Fitting parameters for the empty peak shoulder were obtained from global analysis of empty capsid elution profiles in **(b)**. The remaining portion of the chromatograms was fitted using 5 Gaussian peaks (Fit Peaks 2-6) without parameter constraints to produce a cumulative model with a satisfactory fit quality (adjusted $R^2 > 0.99$). Based on the calculated peak areas, heavy AAV5 sample contains 10.6 ± 0.6 % of light capsids, 84.6 ± 0.8% of heavy capsids, and 4.7 ± 0.2% of the material eluted during a high salt wash with 1M sodium acetate, 20mM BTP pH8.0 (strip peak). Models generated using a different number of fitted peaks with/without Gaussian peak constraints are reported in the table of this section.

### S15. Size Exclusion Chromatography (SEC) for characterizing AAV

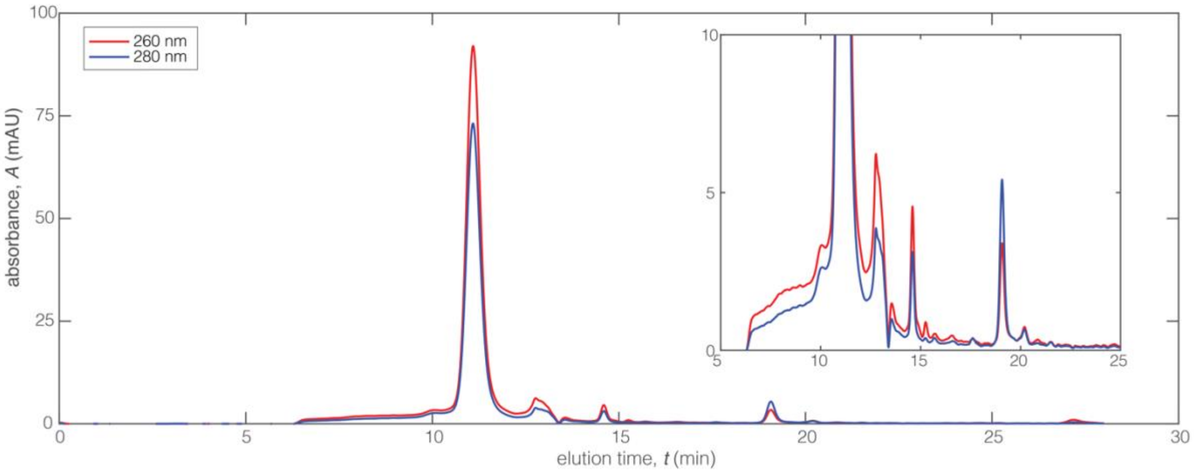


**Figure S15 | Size Exclusion Chromatography (SEC)**. Absorbance spectra of SEC column for two wavelengths UV at 260 nm (red) and 280 nm (blue) vs elution time. Inset shows zoomed-in absorbance spectra (same units) for elution times $5 \leq t \leq 25$ min The 'tailing' at lower elution times indicates presence of aggregates, which being bigger than single AAV particles, elute before than the latter (elution time $t < 10$ sec). Elution time depends on size, the method does not separate heavy to light capsids. However, the peak at $t \cong 11$ sec indicates that the majority of the capsids are heavy since absorbance at 260 nm is higher than 280 nm.
